# *BigNeuron*: A resource to benchmark and predict best-performing algorithms for automated reconstruction of neuronal morphology

**DOI:** 10.1101/2022.05.10.491406

**Authors:** Linus Manubens-Gil, Zhi Zhou, Hanbo Chen, Arvind Ramanathan, Xiaoxiao Liu, Yufeng Liu, Alessandro Bria, Todd Gillette, Zongcai Ruan, Jian Yang, Miroslav Radojević, Ting Zhao, Li Cheng, Lei Qu, Siqi Liu, Kristofer E. Bouchard, Lin Gu, Weidong Cai, Shuiwang Ji, Badrinath Roysam, Ching-Wei Wang, Hongchuan Yu, Amos Sironi, Daniel Maxim Iascone, Jie Zhou, Erhan Bas, Eduardo Conde-Sousa, Paulo Aguiar, Xiang Li, Yujie Li, Sumit Nanda, Yuan Wang, Leila Muresan, Pascal Fua, Bing Ye, Hai-yan He, Jochen F. Staiger, Manuel Peter, Daniel N. Cox, Michel Simonneau, Marcel Oberlaender, Gregory Jefferis, Kei Ito, Paloma Gonzalez-Bellido, Jinhyun Kim, Edwin Rubel, Hollis T. Cline, Hongkui Zeng, Aljoscha Nern, Ann-Shyn Chiang, Jianhua Yao, Jane Roskams, Rick Livesey, Janine Stevens, Tianming Liu, Chinh Dang, Yike Guo, Ning Zhong, Georgia Tourassi, Sean Hill, Michael Hawrylycz, Christof Koch, Erik Meijering, Giorgio A. Ascoli, Hanchuan Peng

## Abstract

*BigNeuron* is an open community bench-testing platform combining the expertise of neuroscientists and computer scientists toward the goal of setting open standards for accurate and fast automatic neuron reconstruction. The project gathered a diverse set of image volumes across several species representative of the data obtained in most neuroscience laboratories interested in neuron reconstruction. Here we report generated gold standard manual annotations for a selected subset of the available imaging datasets and quantified reconstruction quality for 35 automatic reconstruction algorithms. Together with image quality features, the data were pooled in an interactive web application that allows users and developers to perform principal component analysis, *t*-distributed stochastic neighbor embedding, correlation and clustering, visualization of imaging and reconstruction data, and benchmarking of automatic reconstruction algorithms in user-defined data subsets. Our results show that image quality metrics explain most of the variance in the data, followed by neuromorphological features related to neuron size. By benchmarking automatic reconstruction algorithms, we observed that diverse algorithms can provide complementary information toward obtaining accurate results and developed a novel algorithm to iteratively combine methods and generate consensus reconstructions. The consensus trees obtained provide estimates of the neuron structure ground truth that typically outperform single algorithms. Finally, to aid users in predicting the most accurate automatic reconstruction results without manual annotations for comparison, we used support vector machine regression to predict reconstruction quality given an image volume and a set of automatic reconstructions.

## INTRODUCTION

Quantification of neuronal morphology is an essential process in defining neuronal types, assessing neuronal changes in development and aging, determining effects of brain disorders and treatments, and providing important parameters for neuronal computations. However, quantifying the three-dimensional structure of neuronal trees has remained a long-standing challenge in neuroscience. Reconstructing neural structure is essential for understanding cellular scale connectivity and electrophysiological behavior within neuronal networks. Neuron reconstructions have become an essential tool for studying brain circuits and exploring brain function in computational neuroscience (Meijering 2010; Parekh and Ascoli 2013). Neuroscientists and computer scientists alike have been developing and exploring methods for fully automated neuron reconstruction for nearly four decades (Capowski 1983; Senft 2011). While automatic reconstruction of neuron tree structures based on 3D microscopy imaging datasets was expected to be a feasible, if not easy, task for computers, experience during the last decades has underlined the difficulty of this challenge. Diversity among animal species, developmental stages, brain location, and image quality of microscopy datasets implies that algorithms with an impressive performance in small sets of images do not generalize well when applied to image volumes obtained by different research teams.

Recent technological advances in labeling (Cai et al. 2013; Nern, Pfeiffer, and Rubin 2015; Daigle et al. 2018), tissue preparation (Chung et al. 2013; Hama et al. 2011), and imaging techniques (Huisken et al. 2004; Verveer et al. 2007; A. Li et al. 2010) allow both small neuroscience laboratories and especially-large-scale brain science projects (Chiang et al. 2011; Meissner et al. 2020; Markram 2006; Ecker et al. 2017; Stockton and Santamaria 2017) to generate increasingly large fluorescence microscopy datasets for the reconstruction of single neurons. To this end, several automatic reconstruction algorithms have been proposed in the last decades (Wang et al. 2011; Xiao and Peng 2013; Santamaría-Pang et al. 2015; Peng et al. 2017), and individual groups have tackled the challenge of applying automatic neuron reconstruction by mainly focusing on their own datasets (Winnubst et al. 2019; Peng et al. 2021). Even though further improving labeling and imaging quality is key for simplifying the task of automatic reconstruction (Zhong et al. 2021), manual correction and fine-tuning of the automatic results by experts are still needed, being a highly demanding bottleneck for the throughput of those projects. A faithful annotation of the fine details of neuron morphology is particularly relevant for estimating potential connectivity, given that artifacts introduce biases in the estimation of connections between brain regions. This is even more relevant when whole brain modeling techniques rely on synthetic generation of neuronal populations based on available annotation data (Shillcock et al. 2016). Similarly, fine-scale neuromorphological properties such as the radius of the neurite segments have a crucial impact on the electrical modeling of neurons, and seemingly small artifacts in the reconstruction process can lead to fully altered tree topology leading to unreliable simulation of signal integration and transmission. Thus, leveraging our understanding of the performance of the available algorithms and how they match with specific characteristics of different imaging datasets is an essential step toward reaching fully automatic neuron reconstruction (Peng et al. 2015).

Two common problems identified by neuroscientists interested in using reconstruction algorithms are that (1) imaging quality varies considerably among labeling and imaging techniques in different laboratories, and (2) the growing list of available algorithms complicates testing their suitability for specific tasks in a systematic and fast manner. Similarly, algorithm developers lack a standard set of images for benchmarking. The DIADEM challenge (https://diadem.janelia.org/history.html, Gillette, Brown, and Ascoli 2011) is an example of a successful initiative toward standardized benchmarking. However, the diversity of datasets tested in the challenge was relatively limited for studying the relevance of image quality features, and, since this challenge, several novel algorithms have been developed (Peng, et al, 2015). Thus, an interactive extendable framework for benchmarking existing and novel tools in diverse datasets was missing.

The *BigNeuron* project was devised to address these challenges and advance toward a consensus on how to use and improve automatic neuron reconstruction tools. The results presented here summarize the goals reached through its completion, including (1) gathering and sharing a community-contributed, diverse and extensive set of 3D neuron imaging datasets, (2) providing gold standard annotations for a selected subset of images to be used as reference for bench testing, (3) organizing collaborative events for the development of novel automatic reconstruction algorithms, (4) providing a platform for benchmarking existing and novel algorithms against gold standard reconstructions, (5) integrating the obtained knowledge to improve the accessibility, accuracy, and efficiency of automatic reconstruction methods, and (6) providing a tool to suggest the best automatic reconstruction algorithm in external datasets based on our results.

## RESULTS

### An open bench-testing platform for neuron reconstruction

We aimed to bench-test on a common open platform open-source, automated neuron reconstruction algorithms using large-scale, publicly available single 3D neuron image datasets acquired by diverse light microscopy systems. A total of 11 community hackathons and events were held in the first phase.

The *BigNeuron* project utilizes neuron image stacks from different species (including fruitfly and other insects, fish, turtle, chicken, mouse, rat, and human) and nervous system regions such as cortical and subcortical areas, retina, and peripheral nervous system. The data includes multiple light microscopy modalities, especially laser scanning microscopy (confocal/2-photon) and brightfield or epi-fluorescent imaging. The neurons are labeled using different methods, such as genetic labeling and virus/dye/biocytin injection and span a broad range of types (e.g., unipolar, multipolar, releasing different neurotransmitters, and with a wide variety of electrophysiological properties). Many of these image volumes were generated by large-scale neuroinformatics projects such as the Allen Mouse and Human Cell Types projects (http://celltypes.brain-map.org/), Taiwan FlyCircuits (15,921 image volumes; http://www.flycircuit.tw/; Chiang et al. 2011), and Janelia Fly Light (13,449 image volumes; https://www.janelia.org/project-team/flylight), but several data sets are also contributed directly by neuroscientists worldwide (Dataset gathering). In total, about 30,000 single-neuron 3D image volumes were gathered. To generate a representative dataset of various organisms, cell types, and imaging conditions, we randomly selected a small subset from the datasets of the large-scale projects. Thus, 166 neurons were selected for the generation of a diverse set of manually curated “gold standard” reconstructions in annotation workshops and for posterior bench-testing of automatic reconstruction algorithms, defining the Gold166 dataset.

As a result of the hackathons, 16 novel automatic reconstruction algorithms were developed and included in the large-scale bench-testing phase. Some of the novel algorithms have been shown to be suitable for specific practical situations, and 13 of those were published in peer-reviewed scientific journals (Supplementary Materials Table 2). In the bench-testing phase, all ported algorithms were tested on the Gold166 dataset primarily using TITAN at Oak Ridge National Laboratory (USA), as well as supercomputers at Lawrence Berkeley National Laboratory (USA) and Human Brain Project (Europe).

The community effort (Fig. 1) of multiple institutions and single research labs involved in the project yielded several notable outcomes:

- In a series of hackathons worldwide, developers learned (from each other) the relative pros and cons of various methods and how to leverage existing resources to refine or develop new algorithms.
- The project has served as a practical guideline for neurobiologists in determining the suitability of specific reconstruction methods for a variety of image datasets and providing feedback regarding the utility of various sample preparation and imaging protocols.
- Bringing neuron reconstruction methods and results together also encouraged method developers to collaborate, share, and reuse each other’s software modules.
- We developed one of the largest community-derived phenotype databases for single neurons, cataloging neuron shape and projection patterns from different species and different brain regions, and offering an opportunity to mine and query the patterns of neurons with distinct shapes.

**Figure 1.**
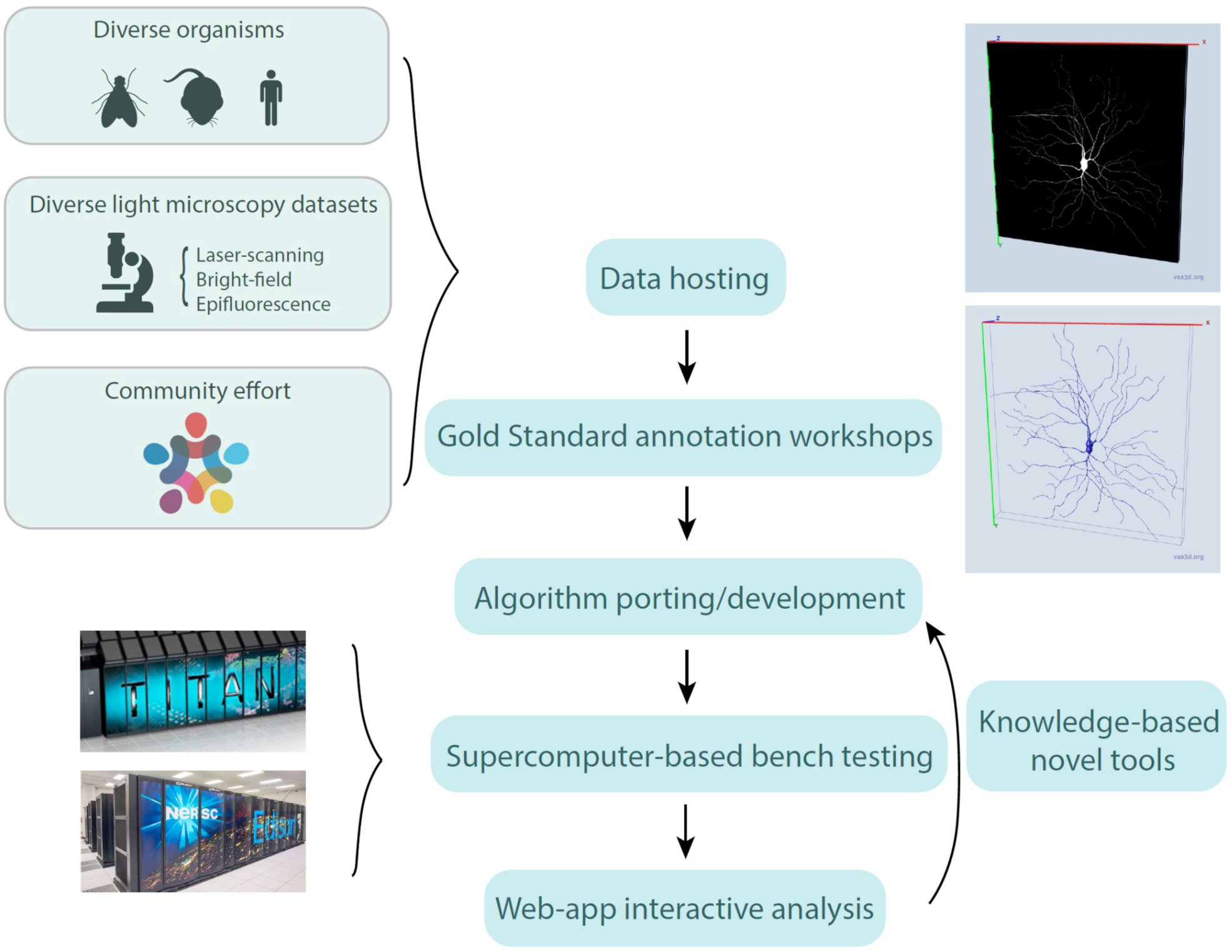
Collaborative efforts led to an open bench-testing platform for neuron reconstruction. In neuroscience labs worldwide, neuronal morphology image stacks are acquired through multiple microscopy modalities from a diverse array of experimental preparations. At the same time, a broad range of reconstruction algorithms is continuously developed using various programming languages, computational platforms, and custom testbeds. Harnessing powerful supercomputers and the open-source community, *BigNeuron* brings together tens of thousands of state-of-the-art 3D neuronal images with all major classes of automated tracing systems to generate an unprecedented number of digital morphological reconstructions. Expert cross-analysis of the initial bench testing dataset against a manually curated gold standard allowed us to generate a unique toolbox of morphometric measurements. Using this knowledge, we developed (1) an interactive web app that allows any user to find similarities between their data and the *BigNeuron* dataset, (2) a method for obtaining consensus reconstructions, and (3) a tool for neuroscientists to predict the best automatic reconstruction algorithm in any single neuron imaging dataset. Adapted from (Peng et al., 2015).

### A web app to allow interactive navigation of heterogeneous bench-testing results

The data generated through the bench-testing phase has produced an extensive collection of heterogeneous results. To allow user-defined interactive exploration of the data, we organized gold standard annotations, automatic reconstructions, their imaging datasets, and associated metadata in R data frames. For bench-testing, we measured the distance between all the automatic reconstructions and the respective gold standard annotations, and computed image quality metrics for each dataset. All the data were pooled in an interactive web-app (Shiny: https://linusmg.shinyapps.io/BigNeuron_Gold166 and https://neuroxiv.net/bigneuron/, Fig. 2) that allows users to perform principal component analysis, t-distributed stochastic neighbor embedding (t-SNE), correlation and clustering, visualization of imaging data and reconstruction in 2D projections, and benchmarking of automatic reconstruction algorithms in user-defined data subsets.

**Figure 2.**
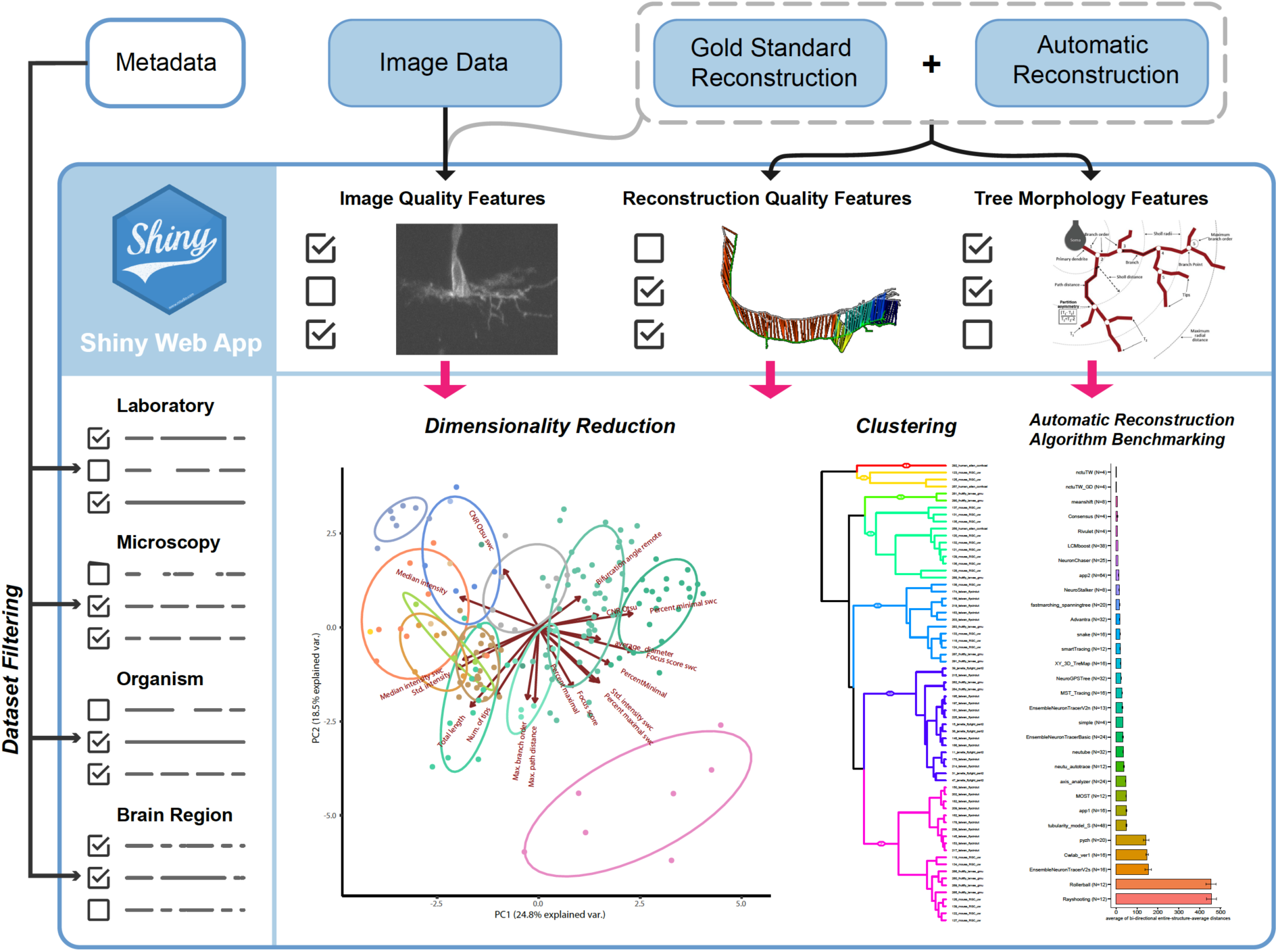
A web app to allow interactive navigation of heterogeneous bench-testing results. Visualization of the Shiny interactive web app (https://linusmg.shinyapps.io/BigNeuron_Gold166/ and https://neuroxiv.net/bigneuron/). The data loaded into the app includes the dataset images, gold standard annotations, automatic reconstructions, and metadata associated with each dataset. Users can interactively choose the image quality and tree morphology metrics used for dimensionality reduction and cluster analysis, and perform reconstruction quality benchmarking.

To encourage developers to continue further automatic reconstruction algorithm development efforts and to simplify benchmarking of novel algorithms, we added to the Shiny app the possibility of uploading and interactively bench-testing reconstruction results of user-defined algorithms. The gold standard preprocessed imaging datasets can be downloaded from https://github.com/BigNeuron/Data/releases/tag/Gold166_v1. After generating single-cell reconstructions for any subset of the data, the obtained automatic reconstructions can be uploaded by specifying the dataset identity (id) of each reconstruction in the filename (see id lookup table https://github.com/lmanubens/BigNeuron/blob/main/lookup_gold166.csv). Once uploaded, the user needs to name the novel reconstruction algorithm and include it in the interactive analysis and benchmarking. We invite developers and users to send novel algorithm implementations and imaging datasets for inclusion in the platform. All submissions will be assessed once per year and introduced in the platform to ensure maintenance of the platform. Reconstruction algorithm implementations and datasets eligible for inclusion in the platform should conform with the guidelines described in (Image quality measurement) and the Open Data Agreement (Open Data Agreement) of the *BigNeuron* project.

### Variance among reconstructions is explained by both image quality and tree morphology features

An overview of the morphological features of the analyzed trees shows that there is high morphological diversity in the analyzed neurons. To quantify the heterogeneity of the data, we computed the coefficient of variation (CV) for a set of morphological features that was also explored in a more extensive dataset (5,099 neurons) in a published neuron reconstruction data mining work using a subset of the reconstructions publicly available at NeuroMorpho.org (Polavaram et al. 2014). For 7 out of 9 neuromorphological features shared among both datasets, the CV of the Gold166 dataset was similar to or higher than its counterpart in the NeuroMorpho.org dataset (Fig. 3A), indicating that the diversity of the Gold166 dataset is sufficient to sample the performance of automatic tracing algorithms in heterogeneous neuron types.

**Figure 3.**
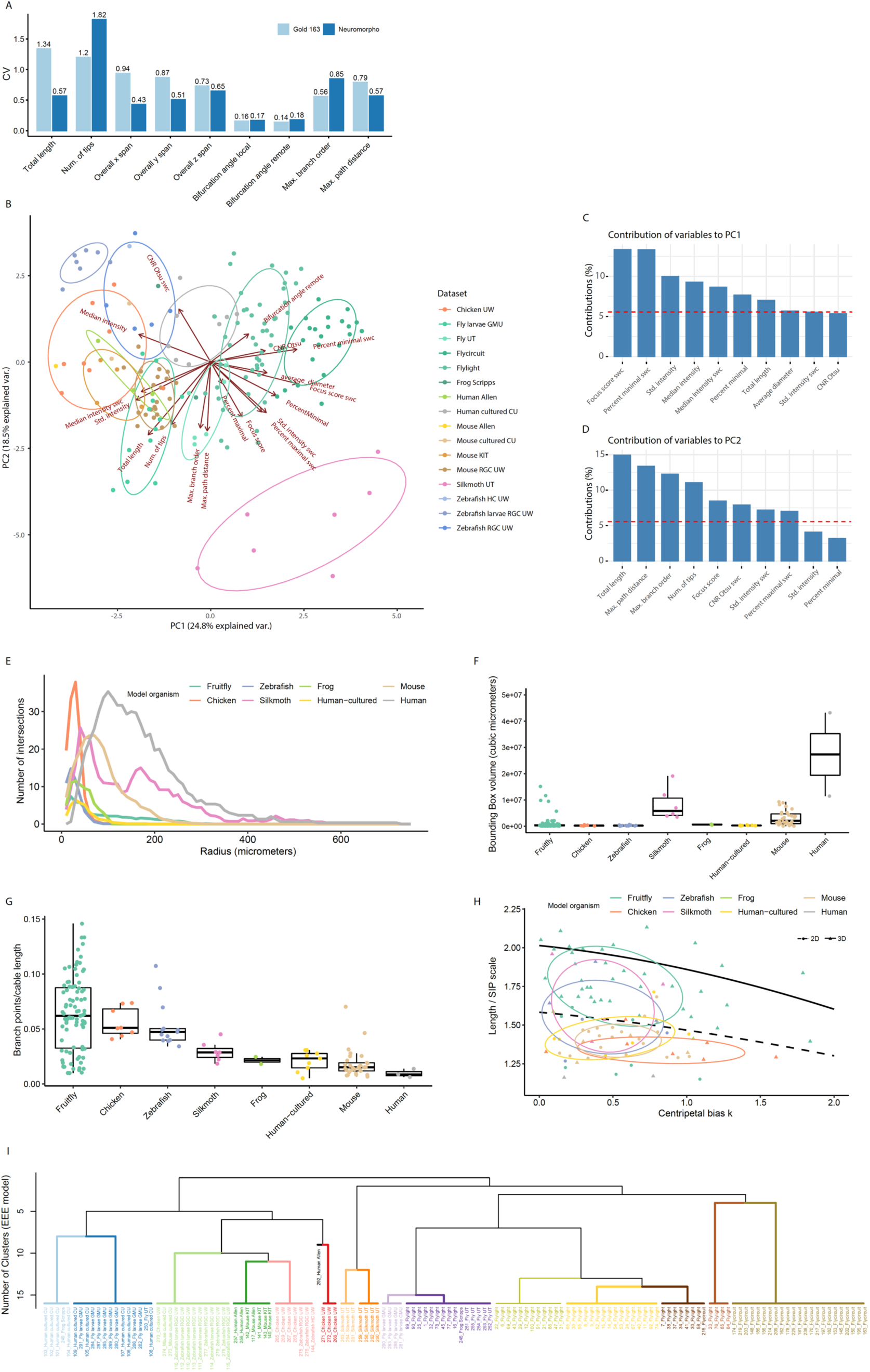
Variance among the datasets is explained by both image quality and tree morphology features. **A** Bar plot showing the coefficient of variation (CV) of various morphological features obtained in the Gold166 dataset, and a dataset with 5,099 neurons previously obtained and analyzed (Polavaram et al., 2014). **B** Principal Component Analysis (PCA) of gold standard neuron reconstructions and their image stack quality metrics. Each point is one gold standard annotation, and the color indicates the dataset it comes from. Arrows represent the direction of each variable in the PCA space. Longer arrows belong to variables that are well represented by the two principal components. Given that 68% of the density of multi variate normal distributions are within 1 Mehalanobis distance of the mean, 68% confidence normal data ellipses for each group are drawn with solid lines. **C-D** Bar plots show the percentage of explained variance for each principal component. Bars represent the contribution (%) of each variable and the red dashed lines indicate the expected average contribution. The first principal component is a composite variable accounting for focus score and the percentage of voxels with minimum intensity in SWC nodes. The second principal component is a composite variable accounting mainly for tree morphology metrics: the total length, maximum path distance, maximum branch order, and the number of tips. **E** Sholl analysis of the neurons in the Gold166 dataset. Each line expresses the average number of intersections quantified at a given distance from the soma for the neurons of a given model organism (color-coded). Neurons from different model organisms present a high diversity of branching patterns. **F-G** Box plots and dot plots show the bounding box volume (**F**) and density of branch points per unit of neurite length (**G**) of the reconstructed cells for each model organism (color-coded). Center line is the median value, the box is limited by Q1 and Q3, and the whiskers add 1.5x interquartile range. Colored dot points are overlayed for all measures. **H** Proportionality constant between the total length and Sholl Intersection Profile (SIP) as a function of the centripetal bias k for planar 2D neurons (dashed black line) and 3D neurons (solid black line). Experimental values for each of the cells in the Gold166 dataset are shown as solid dots color-coded by model organisms. 2D trees are shown as circles and 3D trees as triangles. **I** Hierarchical clustering using both neuromorphological and image quality features. Colors indicate different clusters obtained with a Gaussian Mixture ellipsoidal, equal volume, shape, and orientation (EEE) model (see Supplementary Materials and Fig. S2). Labels indicate the neuron dataset id and the organism they belong to.

To identify the features that account for the variance among datasets, we performed dimensionality reduction using Principal Component Analysis (PCA). The first two principal components explain 43.3% of the variance in the data (Fig. 3B). Principal Component 1 (PC1, 24.8% explained variance) is mainly contributed by measures that account for image quality: the focus score computed on the SWC nodes, the percentage of minimal intensity voxels in the SWC nodes, and the standard deviation of the intensity in the image volume (Fig. 3C). Following those metrics, the median intensity in the image volume, and the median intensity on the SWC nodes show considerable contributions to PC1. Principal Component 2 (PC2, 18.5% explained variance) is mainly contributed by neuromorphological metrics: the total length, followed by the maximum path distance, the maximum branch order, and the number of tips of the trees (Fig. 3D). The focus score and CNR in SWC nodes contribute to PC2 to a lower extent.

A 2D projection overview of those principal components shows that the datasets are clustered by the labs that provided them, being separated along both PC1 and PC2. The variables contributing more to the difference between datasets are mostly related to image quality features, while tree morphology-related features explain most of the variance within each dataset.

Even though a systematic comparison between species should be done for as highly similar cell types (beyond the scope of this work), we present here a comparison of broad neuromorphological differences. Human and silkmoth neurons are big and complex, having branches that reach distances above 200 μm from the soma (Fig. 3E). In the case of silkmoth neurons, this is explained by the fact that the reconstructions include long-range projections (Fig. 3E and 3F), while human dendritic trees span significantly more extensively than all other trees analyzed and have a number of branches above all other organisms (including silkmoth). Mouse neurons are similar to human cells: while being smaller and less complex, they have a comparable ratio of branch points per unit of cable length (Fig. 3G) and cluster together with *ex vivo* human neurons in the PCA (Fig. 3B). In contrast, fruitfly, silkmoth, zebrafish and chicken neurons show ratios of branch points per unit of cable length higher than mammal counterparts (Fig. 3G). While frog neurons show values closer to human and mouse counterparts, the number of cells we analyzed is too small to conclude that this result has biological relevance. Fruitfly neurons differ from silkmoth neurons mainly in terms of maximum path distance and maximum branch order, while showing increased average diameter and bifurcation angles in comparison to zebrafish and chicken neurons, the latter having values closer to human, mouse and chicken neurons. Chicken neurons show high dendritic complexity at small radii (Fig. 3E), but they are closer to zebrafish and fruit fly larvae neurons in regard to size and density of branches per unit of dendritic length (Figs. 3F and 3G).

To explore putative functional heterogeneity in the dataset, we quantified the centripetal bias k. When the centripetal bias is 0, neurites are not preferentially radial to the soma, and their angles toward the root of the tree are distributed uniformly. As the centripetal bias k goes to infinity, all branch segments increasingly have radial directions from the soma. The distribution of branch angles to the tree root in function of k can be expressed analytically as a modified von Mises distribution (Forbes, 2011). It has been shown that the Sholl Intersection Profile (SIP) of specific neuron types can be predicted by their span, total length, and centripetal bias; each of those having specific impacts on neuron functionality (Bird, 2019). Bird et al. also demonstrated the impact of centripetal bias on electrotonic compartmentalization. We assessed putative functional heterogeneity by exploring the ratio between those quantities (Fig. 3H). Our analysis shows that the centripetal bias of the Gold166 neurons is constrained to values lower than 2, indicating that the neurons in this set have low electrotonic compartmentalization and long conduction times in comparison, for example, to hippocampal neurons (with k∼7 and k∼12 for CA1 and DG respectively). All 2D cells show a Length/SIP scale ratio consistent with the theoretical 2D von Mises root angle distribution. Interestingly, chicken neurons, despite being 3D, show a dendritic occupancy comparable to 2D trees, explained by their flat morphology. 3D silkmoth and fruitfly neurons show higher dendritic occupancy consistent with the theoretical 3D von Mises root angle distribution. In terms of cross-species comparison, we observed similar distributions in the centripetal bias of all species with a non-significant tendency to higher values in some chicken and fruitfly 3D neurons.

We performed unsupervised hierarchical clustering using both neuromorphological and image quality features. We found that our data contains 16 clusters and is best approximated by an ellipsoidal, equal volume, shape, and orientation (EEE) model (Fig. S2 in Supplementary Materials, maximum Bayesian Information Criterion of 2989.32). The data are mainly clustered by dataset, and when those were obtained from the same organisms, they usually clustered together.

### Consensus trees provide informed best estimates of ground truth neuronal structure

Given the high diversity of existing automatic reconstruction algorithms, it is reasonable to assume that their performance in reconstructing specific tree morphology features may vary. We tested their accuracy by measuring the error between the morphological features in the automatic reconstructions and the gold standard trees. We found that each morphological feature was best estimated by different algorithms (Fig. 4A; Kruskal-Wallis; average contraction H(5)=153.16, p=2e-30; average fragmentation H(5)=339.19, p=3e-70; bifurcation angle remote H(5)=42.43, p=4.8e- 8; maximum branch order H(5)=61.36, p=9.5e-14; maximum path distance H(5)=12.82, p=0.0251; number of tips H(5)=62.20, p=9e-12; overall x span H(5)=21.90, p=6.9e-4; total length H(5)=23.41, p=2.8e-4). These results suggest that a set of diverse algorithms can contribute complementary information to approximate the ground truth with increased precision. To build on this idea, we developed an algorithm for generating “consensus trees” based on a set of automatic reconstructions (see Generation of consensus trees). By iteratively averaging the closest node positions of the set of reconstructions, the algorithm defines a set of consensus nodes (Fig. 4B). We assigned a confidence value to each consensus node by voting their existence in each individual automatic reconstruction. Thus, nodes highly prevalent in many reconstructions are kept in the consensus tree, while false positive fragments in small numbers of reconstructions have low confidence and are discarded. Finally, the high confidence set of consensus nodes obtained through this process is connected in a single tree using the Minimum Spanning Tree algorithm (Fig. 4B and 4C, Generation of consensus trees). To assess the ability of the consensus tree algorithm to provide informed best estimates of gold standard reconstructions, we bench-tested all the automatic reconstruction algorithms by measuring the distance between each automatic reconstruction and their corresponding gold standard tree. The bench testing results showed that, in 5 of 16 clusters identified in the data (Variance among the datasets is explained by both image quality and tree morphology features, Fig. 3I), the consensus tree algorithm provided the most accurate approximation to gold standard trees (Fig. 4C and 4D). However, in the other clusters, different algorithms outperformed the consensus tree strategy (Fig. 4E).

**Figure 4.**
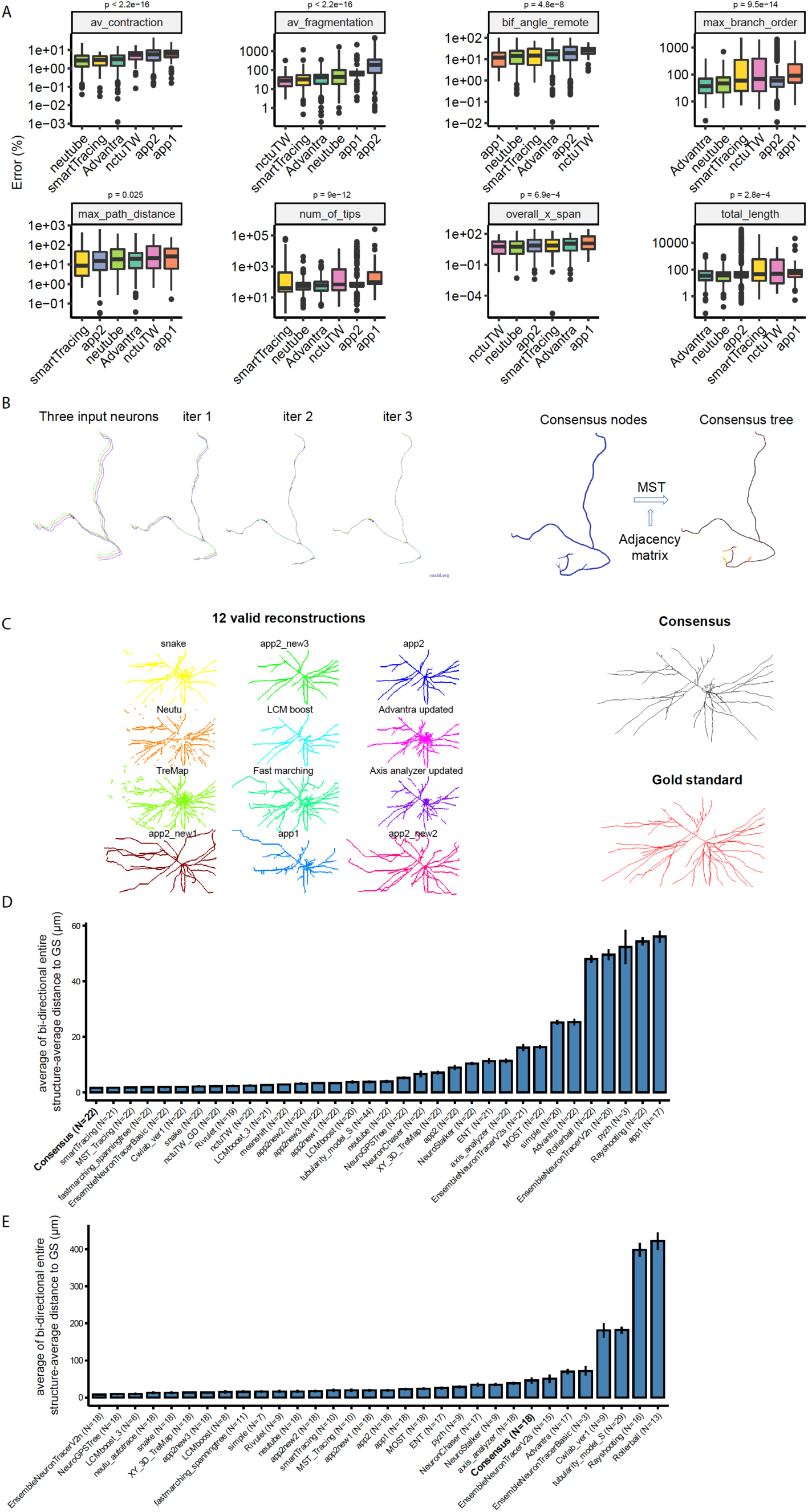
Consensus trees provide informed best estimates of ground truth neuronal structure. **A** Errors for a set of morphological metrics in a random subset of automatic reconstruction algorithms. The error was computed as the difference between the metrics obtained in the gold standards and the automatic reconstructions. Errors for the automatic reconstructions obtained from all image volumes with a given algorithm are shown as boxplots. Center line is the median value, the box is limited by Q1 and Q3, the whiskers add 1.5x interquartile range, and outliers are indicated with points. P-values were obtained with Kruskal-Wallis (one-sided) tests. Numbers of reconstructions for each method: Advantra=109, app1=98, app2=102, nctuTW=54, neutube=112, smartTracing=86. **B** Overview of the development of a novel algorithm to generate consensus trees. The algorithm first performs Iterative Match and Center: for each node in each tree, it identifies the nearest corresponding location among input neurons, and shifts to the mean location. Nodes from all input neurons are merged to form consensus nodes and reliability weights among consensus nodes are established by collecting votes from the connections from individual input neuron trees. A Maximum Spanning Tree algorithm is used to connect consensus nodes to form the consensus tree. **C** An example of the results obtained with the consensus tree algorithm using a set of 12 automatic reconstructions. **D** Bidirectional entire structure average distance between automatic reconstructions and the gold standard of the dataset cluster number 9 (see Fig. 3I). Mean and Standard Errors are shown as a barplot. **E** Bidirectional entire structure average distance between automatic reconstructions and the gold standard of the dataset cluster number 8 (see Fig. 3I). Mean and Standard Errors are shown as a barplot.

### Image quality metrics correlate with algorithm performance

To explore how reconstruction quality varies among different images, we tested how the distance between consensus reconstructions and gold standard trees correlate with every feature of the datasets. We performed this analysis by obtaining a correlation matrix among reconstruction quality, image quality, and neuron morphology features of all the automatic reconstruction results (Fig. 5A). Distance metrics correlated with diverse image quality and tree morphology features: CNR, CNR in SWC nodes, the median intensity of the image, parent-daughter ratio, and remote bifurcation angle (Fig. 5A, cluster 3). The second cluster of features showed a negative correlation with distance metrics: focus score (together with its equivalent obtained in SWC nodes), the percentage of image voxels with minimal intensity (together with its equivalent in SWC nodes), the average diameter of the automatic reconstruction segments and the standard deviation of the image intensity in SWC nodes (Fig. 5A, cluster 1). Finally, a set of features did not show any correlation with the quality of the consensus reconstructions: the number of tips, maximum branch order, total length, maximum path distance, the standard deviation of the image intensity, the median intensity of the image in SWC nodes and number of stems (Fig. 5A, cluster 2). Thus, as a representative example, the percent of different structure between consensus reconstructions and gold standards positively correlates with the image CNR (Otsu) (Fig. 5B, Pearson’s correlation coefficient: R^2^=0.23, p<1.3e-8). By contrast, the percent of different structures negatively correlates with the parent-daughter ratio (average ratio of the number of child nodes for each parent node in the tree structure; Fig. 5C, R^2^=0.2, p=4.1e-8) and also negatively correlates with the average remote bifurcation angle (Fig. 5D, R^2^=0.2, p=7.1e-8), indicating those tree morphology features have high values in low accuracy automatic reconstructions.

**Figure 5.**
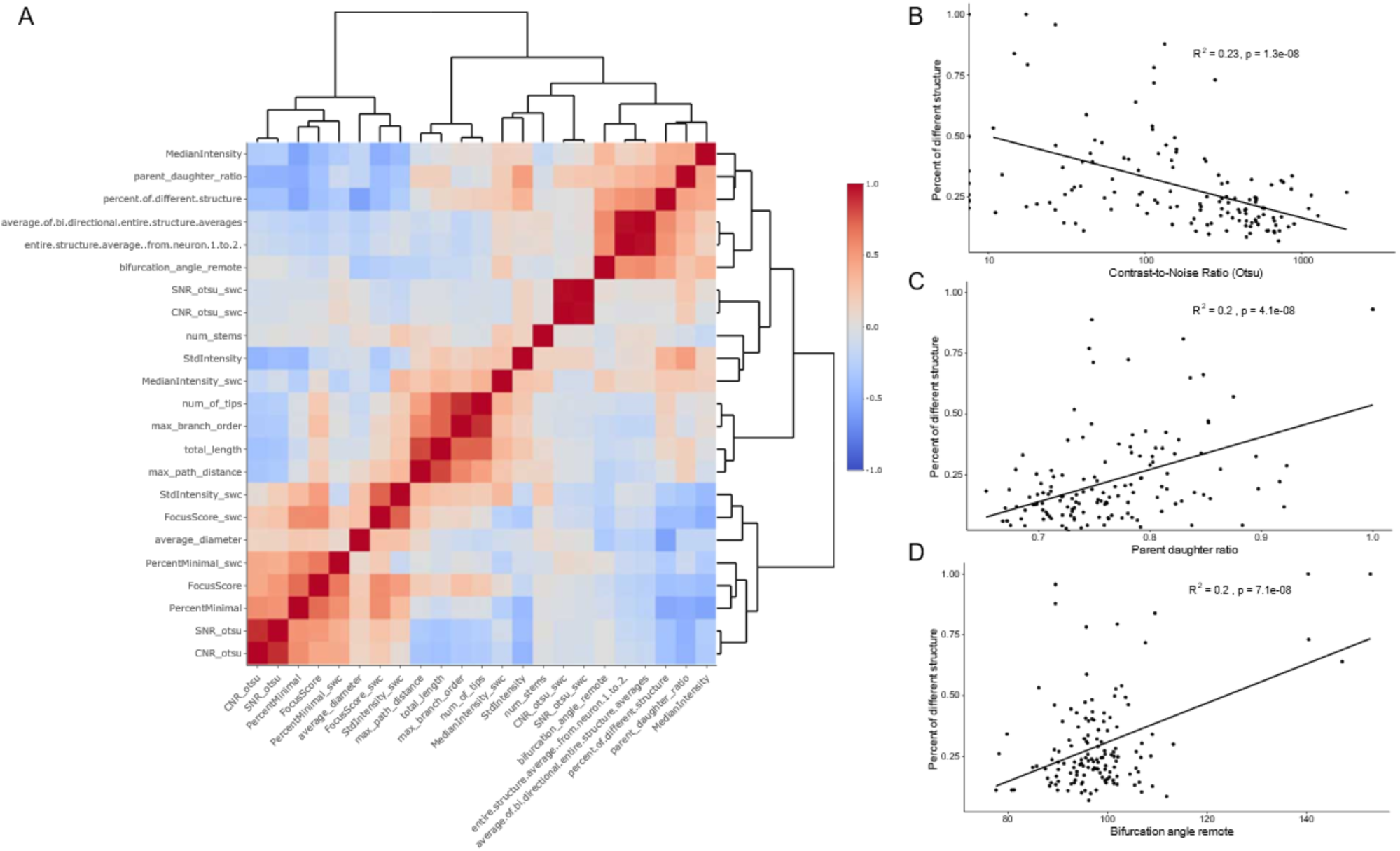
Image quality metrics correlate with best algorithm performance. Hierarchical clustering among image quality metrics, tree morphological features, and reconstruction quality. Reconstruction quality correlates with a set of features, indicating that more focused images of big neurons tend to provide better automatic reconstruction results. **A** The heatmap indicates color-coded pairwise Pearson correlations between metrics obtained for consensus tree reconstructions. **B-D** Correlation plots for image quality and dendritic tree morphology features (**B**: CNR Otsu, **C**: parent-daughter ratio, and **D**: bifurcation angle remote) and consensus reconstruction quality (% of different structure).

### Support vector machine regression for predicting best algorithm performance

These results suggest that specific tree morphology features measured from a set of diverse automatic reconstruction results, together with image quality metrics, can be informative toward choosing the most accurate option within a set of algorithms. To take advantage of this information, we used a Support Vector Machine (SVM) that, given an image volume and a set of automatic reconstructions, predicts the best reconstruction algorithm in the set based on a regression of the percentage of difference between the automatic reconstructions and gold standards (Fig. 6). By training using 85% of our data, we obtained a regression that allows us to predict the percentage of difference between the automatic reconstructions and gold standards (Coefficient of determination of 0.661; Fig. 6A). Learning curves with increasing data percentage used for training show that the regression quality reaches a plateau with slight improvement once more than 30% of the data is used for training (Fig. 6B), suggesting that this strategy will generalize well when it is used in novel neuron reconstruction datasets. We assessed the quality of the predictions by comparing the percentage of different structures of all the automatic reconstructions, the true best algorithms, and the predicted best algorithms for each dataset. Our analysis showed that both known best algorithms and predictions in most of the cases performed better than algorithms chosen by chance (Fig. 6C and 6E; the percentage of difference distribution is positively skewed, medians of 18%, 22%, and 44% for true best, predicted and all algorithms respectively; Wilcoxon test All algorithms vs. True best V=20145, p=5.12e-10, n1=1113, n2=20; All algorithms vs. Predicted best V=16470, p=2.32e-4, n1=1113, n2=20; Predicted best vs. Predicted worst V=19, p=3.03e-8, n1=20, n2=20). Similarly, algorithms predicted to be worse showed a negatively skewed distribution in the percentage of difference to gold standards (Fig. 6C; median of 99%). Interestingly, this analysis highlighted the fact that for a few neuron datasets, none of the automatic reconstruction algorithms was able to recapitulate the gold standard annotations. This was found to be the case, for example, of chicken cells, which showed particularly low signal-to-noise ratio in their neurites and have specific distinctive morphological aspects (such as an increased soma size, high branch density, and high centripetal bias).

**Figure 6.**
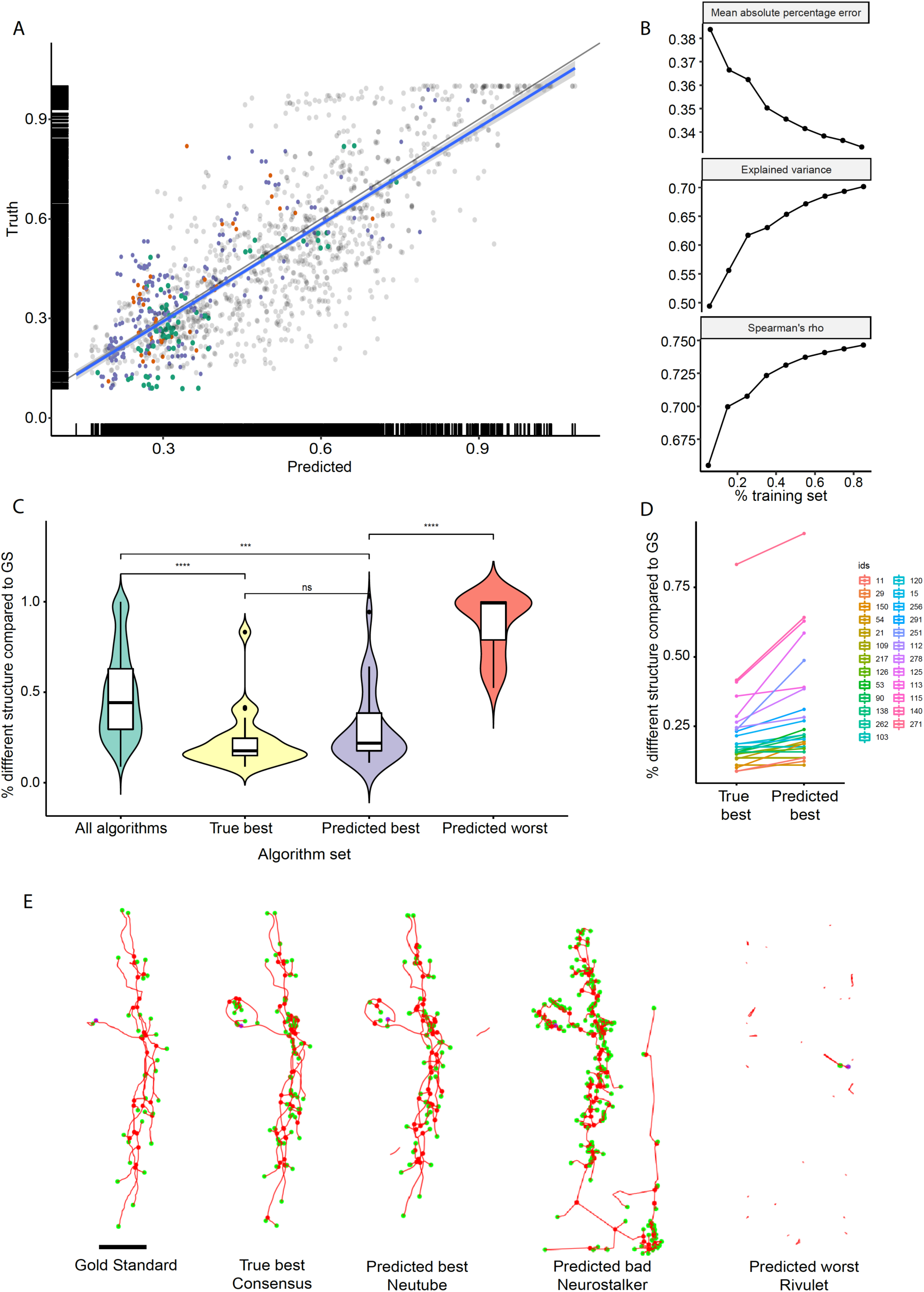
Support vector machine regression for predicting best algorithm performance. **A** Support vector machine regression of the percentage of difference between automatic reconstructions and gold standards. Each point represents an automatic reconstruction and the true and predicted percentage of difference values. Colored points show subsets of results for individual datasets. The rest of the data in the testing set is represented in gray. **B** Learning curves showing the quality of the regression as a function of the percentage of data used for training. Each point is the average of 5 repetitions using different random subsets of the data for training. **C** Percentage of difference between automatic reconstructions and gold standard annotations represented as an overlay of box and violin plots. Statistical comparisons among groups were performed using the Wilcoxon test (two-sided); ^***^ indicates p<0.001, ^****^ indicates p<0.0001. The number of reconstructions in each group are: All algorithms=1113, True best=20, Predicted best=20, and Predicted worst=20. **D** Pair plot showing the percentage of difference between automatic reconstructions and gold standard annotations for the true best algorithms and the predicted best algorithms. Colors indicate different neuron datasets in the testing set. **E** Representative images of the gold standard annotation, the true best automatic reconstruction algorithm, and various predictions using the regression results. Red lines represent neuron tree branches, blue dots indicate root points, red dots indicate roots of the trees, and green dots indicate terminal points. Scale bar = 100µm.

### Showcase of best algorithm prediction in fMOST data

Recent community efforts, like the BRAIN Initiative Cell Census Network (BICCN, https://biccn.org/), are using fluorescence micro-optical sectioning tomography (fMOST) to map neuron architecture within whole mouse brains. To illustrate the value of *BigNeuron* in this community, we used the SVM regression model trained with the Gold166 dataset to predict best-performing algorithms in fMOST image volumes. The SVM model provided predicted values for the percentage of difference to gold standard trees (not available for those images). The algorithms predicted to provide the best results were neutube and Consensus (Fig. 7A and 7E). Visual inspection of the reconstructions suggests that the regression model provides a reasonable approximation toward the best automatic algorithm selection (Fig. 7B-D and 7F-H). A comparison of image quality features between the Gold166 and fMOST images showed that the most similar dataset to fMOST was the Zebrafish larvae RGC cell set (Fig. 7J). The bench-marking of automatic reconstruction algorithms in the images of this set shows that consistently with the predictions, neutube and Consensus algorithms are among the best approximations to gold standard reconstructions.

**Figure 7.**
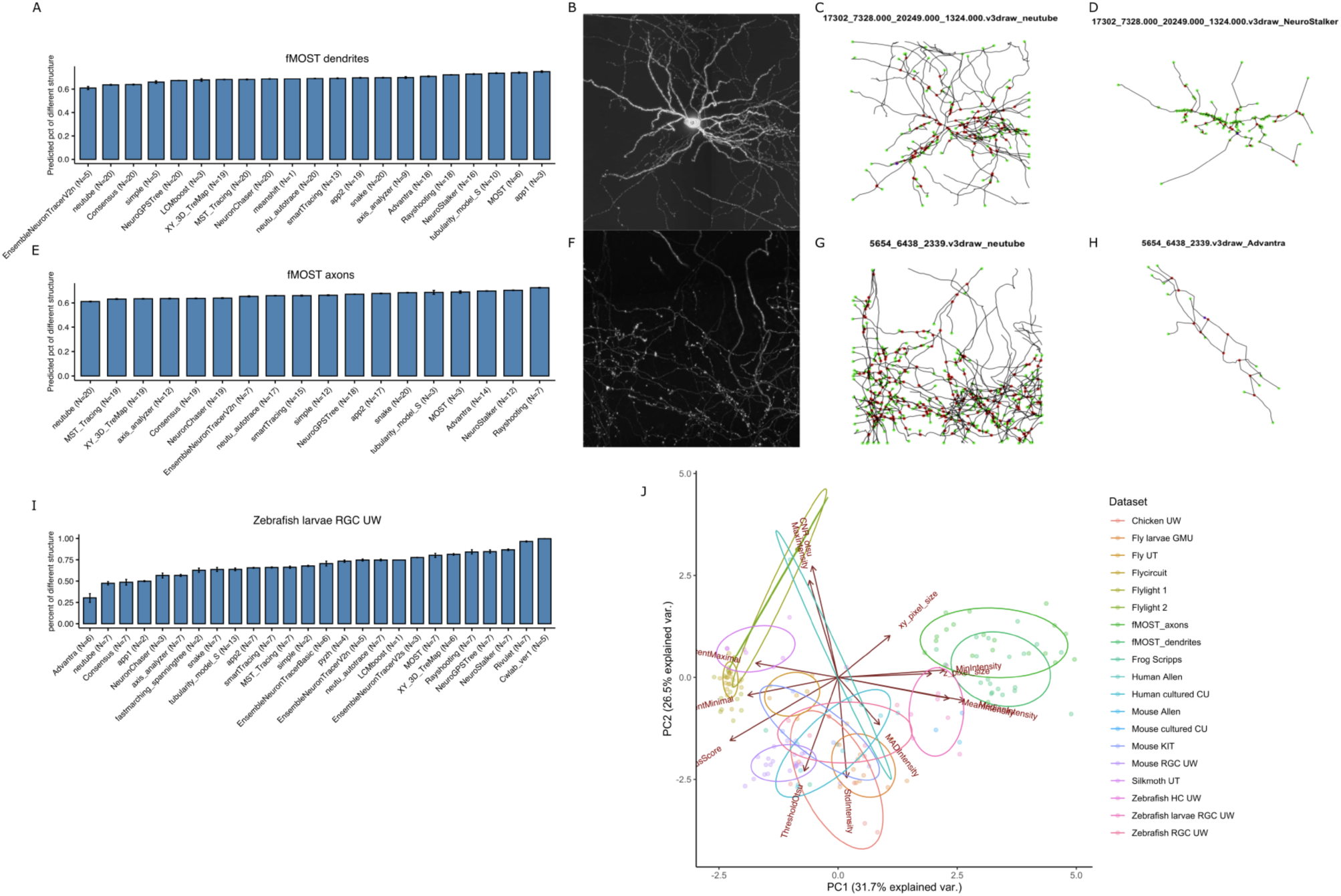
Showcase of best algorithm prediction in fMOST data. Comparison of results obtained for fMOST dendrite and axon image blocks, including the predicted percentage of different structure and representative visualizations. In terms of image quality features, fMOST data is similar to the Zebrafish larvae RGC UW dataset from *BigNeuron* gold standard data, for which bench-marking results are comparable. **A** Predicted percentage of different structure for automatic reconstructions of fMOST dendrite image blocks. **B** Maximum intensity projection of a representative fMOST dendrite image block. **C-D** 2D projections of the neutube (C) and NeuroStalker (D) algorithms automatic reconstructions in the image volume represented in B, predicted to provide best and low accuracy results respectively. **E** Predicted percentage of different structure for automatic reconstructions of fMOST axon image blocks. **F** Maximum intensity projection of a representative fMOST axon image block. **G-H** 2D projections of the neutube (G) and Advantra (H) algorithms automatic reconstructions in the image volume represented in B, predicted to provide best and low accuracy results respectively. **I** Percentage of different structure between automatic reconstructions and gold standard trees for the Zebrafish larvae RGC neurons. Mean and Standard Errors of percentage of different structure are shown as bar plots. **J** Principal Component Analysis of gold standard datasets accounting for their image quality metrics. Each point is one gold standard image volume, and the color indicates the dataset it comes from. Arrows represent the direction of each variable in the PCA space. Longer arrows belong to variables that are well represented by the two principal components. Given that 68% of the density of multi variate normal distributions are within 1 Mehalanobis distance of the mean, 68% confidence normal data ellipses for each group are drawn with solid lines.

## DISCUSSION

The complexity of multi-modal cell type characterization requires reproducible and high-fidelity neuronal reconstructions and precise estimates of their variability. Although quantification of neuronal morphology is an important approach in neuroscience, how to accurately and efficiently perform this task across diverse types of neuronal images in an unbiased fashion has been a long-standing challenge. This challenge has become even more acute with the introduction of exciting new technologies that generate 3D, complete neuronal morphology datasets at high speed (Chung and Deisseroth 2013; Zheng et al. 2013; Verveer et al. 2007; Jiang et al. 2022). Moreover, large-scale brain science projects such as the BRAIN Initiative in the USA (Alivisatos et al., 2012), Europe’s Human Brain Project (Kandel et al., 2013), and the Allen Cell Types Database (http://celltypes.brain-map.org/) integrate reliable automatic reconstruction in their production pipelines to allow fast quantification of the structure of hundreds of thousands of neurons. Motivated by these demands, the *BigNeuron* project has contributed to improving existing automatic reconstruction standards, focusing on the interaction and discussion among neuroscientists and developers, and integrating data generation and development efforts from several laboratories. *BigNeuron* takes an innovative approach toward the standardization and development of new automatic reconstruction methods, including events for discussion and identification of major issues, and coordinated hackathons for the development of tools to overcome them. Data and some results from this effort have been published online for the community to take advantage of them.

The BigNeuron project has inspired a new generation of technical approaches to measure and compare the similarity of neuronal trees (Costa et al. 2016; Kanari et al. 2018). The experience obtained highlighted major challenges in the field of neuron annotation and classification (Meijering et al. 2016; Zeng and Sanes 2017), and contributed to justify new methods for imaging (Treweek et al. 2015; Ke et al. 2016) and large-scale annotation of neurons (Gong et al. 2016; Winnubst et al. 2019; Peng et al. 2021). The project has been presented in 9 scientific meetings, highlighted in 10 press releases, and has led to 11 peer-reviewed publications in scientific journals and 3 data releases (see Supplementary Materials for details). However, a systematic analysis of the tools and results generated by these projects is missing. In this work, we developed the *BigNeuron* interactive analysis web app, a platform for the neuroscience community that allows end-users of automatic reconstruction tools to explore interactively the results obtained with a great diversity of open-source algorithms and to predict which of them will provide the best results in their specific datasets. *BigNeuron* allows developers to interactively benchmark their tools against the methods assessed in this work, and to introduce datasets and novel algorithms within the benchmarking set upon request. This collaborative standardized approach is significant because the problem of choosing a method to perform automatic reconstruction in project-specific datasets is an arduous task. Having a standard resource to compare and analyze the methods’ performance, while introducing tools to quantitatively assess features of the images relevant to the problem, is an approach that can provide easy-to-apply contrastable methods to solve it.

The diversity in the data that we analyzed allowed us to gain relevant insights toward future steps in improving automatic reconstruction tools. We first focused on a major aspect usually overseen when novel reconstruction algorithms are developed and tested: the variance among the datasets obtained by different teams. We found that both neuromorphology and image quality features explain comparable amounts of variance among the analyzed data, image quality being a relevant aspect to take into account in the process of choosing the most suitable tool for a specific dataset. By considering this information, we benchmarked existing and newly developed algorithms for different types of datasets. We found that different types of datasets are best reconstructed by specific algorithms. Also, individual neuromorphological features are best approximated by particular algorithms. Thus, the information obtained through the pooling of datasets and benchmarking of reconstruction tools can be used to generate novel knowledge-based improved algorithms. Previous works have used the *BigNeuron* datasets to benchmark novel algorithms (Li et al. 2017; Liu et al. 2018; Gu et al. 2020; Radojević and Meijering 2019) and to develop neuromorphological features for the analysis of dendritic trees (Kanari et al. 2018). However, to our knowledge there is only one study that explores the idea of using an ensemble learning algorithm to generate improved automatic reconstruction results (Wang et al. 2017), an algorithm included within our analysis. As another step in this direction, we developed a tool to generate consensus trees based on a set of automatic reconstructions. Our results highlight the potential of such an approach, providing the best estimates of the ground truth neuron structure in many of the datasets we analyzed.

While the Ensemble Neuron Tracer and our Consensus Tree algorithms implicitly rely on identifying and discarding unrealistic features in automatically generated trees, our data sets and web app allow exploring interactively the relationship between reconstruction quality and all the computed features of the analyzed trees. Our results point at several image features (e.g. image intensity correlation) as main correlates of reconstruction quality, and we also have identified weak correlations with neuromorphological features such as the parent-daughter ratio. Those observations can further inform future algorithm development by adding morphological constraints to the generated trees. Even though we generated data for a few variations in parameter sets for some of the methods, in this work, we focused on the comparisons among base methods. We think future efforts could focus in the iterative search of optimal parameter sets of specific methods by applying optimization algorithms with maximization of reconstruction quality as the objective function.

Finally, to help end-users identify optimal performing algorithms for their datasets, we used a support vector machine regression to predict the best algorithm performance. We developed a set of Vaa3D plugins that allows users to run all the automated reconstruction algorithms included in *BigNeuron* and quantify the image quality features of their datasets. By performing regression against all the analyzed datasets, users obtain a prediction for the best algorithm without the need of generating a set of gold standard reconstructions for bench-testing. In most cases, our regression method provides accurate estimates of the automatic reconstruction quality. However, there is a small number of datasets for which there is a substantial difference between the true and the predicted best algorithms. Thus, this tool should be used with caution. Interestingly, in a few cases, none of the automatic reconstruction methods was able to produce any result close to the ground truth (>50% of different structure). This happened in datasets with extremely low signal-to-noise ratios and overall bad imaging quality. Quantitative analysis of those features can help users to find objective thresholds for image quality features below which datasets should be discarded before spending time and resources on automatic reconstruction methods. We think it is necessary to explore further refinement of the machine learning predictions of best-performing algorithms introduced in this work. Recent projects are generating unprecedented volumes of gold standard reconstructions (e.g. Peng et al. 2021). We expect that applying the methods presented in standardized, larger gold standard datasets will provide a valuable tool toward full automation of neuron reconstruction in specific combinations of microscopic imaging and labeling techniques that are becoming standards in the field of structural interrogation of single neurons in the whole mouse brain.

## Data availability

3D image volumes of the Gold166 dataset and gold standard reconstructions are available at http://web.bii.a-star.edu.sg/bigneuron/gold166.zip. Bench-testing automated reconstructions can be downloaded from https://github.com/BigNeuron/Data/releases/tag/gold166_bt_v1.0. The complete set of image volumes gathered throughout the project, amounting to ∼4TB of data, is available upon request.

## Code availability

The source code developed is released open source and is available at https://github.com/lmanubens/BigNeuron. The Shiny web app can be used at https://linusmg.shinyapps.io/BigNeuron_Gold166/ and https://neuroxiv.net/bigneuron/. Source code utomated reconstruction algorithms developed throughout the project can be found at https://github.com/Vaa3D/vaa3d_tools/tree/master/released_plugins/v3d_plugins.

### Acknowledgements

This project was supported by Allen Institute for initialization and a series of events, and a DOE’s Oak Ridge National Lab Leadership Computing award to Hanchuan Peng; the cross-platform bench-test was also supported intensively at Lawrence Berkeley National Lab, Allen Institute for Brain Science, and Blue Brain Project at EPFL, Southeast University at Nanjing, and various other facilities. This project also used large-scale display wall and immersive VR facilities at Imperial College of London, Oak Ridge National Lab, Janelia Research Campus of HHMI, and Southeast University. This project was also supported by Tencent Inc. and Southeast University for interactive online analysis. This project has inspired more than 260 other publications to date. We acknowledge Prabhat, Zhijiang Wan, Jinzhu Yang, Hang Zhou, Arunachalam Narayanaswamy, Shaoqun Zeng, Przemyslaw Glowacki, Dezhe Jin, Zhihao Zheng, Pengyu Hong, Tao Zeng, Rongjian Li, Hidetoshi Ikeno, Yu-Tai Ching, Tingwei Quan, Jan-Felix Evers, Chloe Murtin, Shan Gao, Yan Zhu, Yang Yang, Hiroyuki Ai, Shin Ishii, Edward Hottendorf, Takashi Kawase, and a number of other colleagues for assistance in developing and porting automated reconstruction algorithms. We thank Staci Sorensen for assistance in discussing and tracing some gold standard neuron reconstructions, and Ryohei Kanzaki, Daisuke Miyamoto, Rachel Wong, Yun Wang, Ed Lein, Cher Bass, Steve Danzer and many other colleagues for providing neuron image datasets. We acknowledge Allan Jones for giving the project the name and support throughout the project. We thank Rafael Yuste and David Van Essen for discussion; John Isaac and Kevin Moses for support from Wellcome Trust to organize the Cambridge University hackathon; Gerald Rubin and Nelson Spruston for support, data sharing and event organization at the Janelia Research Campus of HHMI; INCF for financial support for organization of a series of meetings; Beijing University of Technology, Imperial College of London, and Southeast University for support of hackathons and workshops. We thank Xuan Zhao for assistance in hosting the shiny app in the neuroXiv website; Penghao Qian, Zuohan Zhao and Xin Chen for assistance in formatting the manuscript. We thank Anne Carpenter for discussion in organizing a tracing hackathon. We acknowledge the National Science and Technology Innovation 2030 - “Brain Science and Brain-Inspired Research” Program of China (Grant 2021ZD0204002). Bing Ye was supported by the NIH grant R01 EB028159. Ann-Shyn Chiang was supported by the Higher Education Sprout Project co-funded by the Ministry of Education and the Ministry of Science and Technology in Taiwan. We acknowledge National Center for High-Performance Computing in Taiwan for managing the FlyCircuit data.

## Author Contributions

H.P. conceived and conceptualized this project. H.P., G.A., E.M. formed the managing team and communicated with other key investigators including A.R., J.Y., J.R., K.I., J.K., G.J., P.GB., Y.G., N.Z., G.T., S.H., M.H. and C.K. to secure key resources to execute the project. H.P. envisioned and led the development of the computational and data analysis platform. L.MG. developed the data analysis for this study. H.C. assisted in hosting the shiny app in the neuroXiv website. Z.Z. along with the help of H.P. bench-tested neuron reconstruction algorithms and generated the automatic morphology reconstructions. Y.L. bench-tested neuron reconstruction algorithms in fMOST datasets. H.C. and H.P. color-separated single neurons from fruitfly brain images generated by A.N. and colleagues in Gerald Rubin lab and the Janelia FlyLight Project Team. H.C., S.N., A.N., and C.D. participated in tracing hackathons and contributed to the generation of gold-standard test dataset. Y.W., E.R., L.M., B.Y., H.Z., H.T.C., AS.C., J.F.S., M.P., R.L., D.N.C., HY.H, M.S., K.I., J.K., G.J., P.GB., and M.O. contributed testing neuron images. A.B., T.G., Z.R., X.L., Y.L., K.B., L.G., L.C., J.Y., L.Q., S.L., H.Y., W.C., S.J., B.R., CW.W., A.S., P.F., M.R., T.Z., D.I., J.Z., T.L., E.B., E.CS., P.A. and J.S. contributed reconstruction algorithms. L.MG., G.A., E.M., and H.P. wrote the manuscript with assistance from co-authors.

## METHOD DETAILS

### Dataset gathering

14 neuroscience research labs and institutions worldwide acquired the imaging datasets used in this work, combining different imaging methods, model organisms, and neuron types. Table 1 in the Supplementary Materials summarizes metadata describing the relevant aspects of each dataset. All the datasets analyzed in this manuscript are freely accessible at https://github.com/BigNeuron/Data/releases. The complete set of image volumes gathered throughout the project, amounting to ∼4TB of data, is available upon request.

## QUANTIFICATION AND STATISTICAL ANALYSIS

### Image quality measurement

To quantify image quality, we implemented a plugin in Vaa3D (RRID:SCR_002609, version 3.497, http://vaa3d.org) that computes the features described in detail in (Bray and Carpenter 2018). A brief description of the features measured with the plugin can be found in the Supplementary Materials. The Vaa3D plugin used is available at: https://github.com/Vaa3D/vaa3d_tools/tree/master/hackathon/linus/image_quality. To obtain image quality measurements associated with each individual reconstruction, we also computed these features within the 3D data only in the image voxels belonging to SWC nodes of the trees. Specifically, we extended the “image_quality” plugin, allowing us to specify a SWC file associated with an image volume and compute the image quality metrics only using the intensity information of those voxels.

### Image preprocessing

Due to varying properties of image acquisition pipelines among institutions, it is challenging to define a universal data preprocessing protocol for reconstruction. We performed Brainbow color separation, 8-bit data type conversion and color inversion for brightfield images as preprocessing tasks prior to automatic reconstruction algorithm bench-testing, using Vaa3D and its plugins. A brief description of the preprocessing steps can be found in the Supplementary Materials. Figure S1 shows examples of the effect of each preprocessing step in the imaging datasets.

### Generation of gold standard reconstructions

A total of 14 labs contributed 17 datasets spanning a broad diversity of species, brain regions, neuron types, labeling methods, microscopy techniques, and imaging resolution (Supplementary Materias Table 1). Gold standard annotations were produced in the *BigNeuron* Neuron Annotation Workshop at Allen Institute, Seattle, June 15-17, 2015. Each reconstruction entry was manually validated by at least 6 annotators working collaboratively. Notably, due to limited image quality, some reconstructions can be ambiguous. Annotators were asked to make their best judgment in such cases and vote to maximize the agreement between them. Thus, we are confident the final reconstructions reflect the best possible reconstruction generated by human experts from the respective image given the limited image quality, time, and resources; we thus adopted this dataset as the gold standard for the automated reconstruction algorithms. After manual curation and postprocessing, we obtained a gold standard set of 166 reconstructed neurons.

### Development of automatic reconstruction algorithms

We ported 44 implementations of 32 methods for automatic tracing as *BigNeuron* plugins to Vaa3D (Supplementary Materials Table 2), including 16 already existing tracing algorithms and 16 novel ones specifically developed within the scope of this project and not previously published (for these latter ones, we provide a brief description in the Supplementary Materials). Of the 44 implementations, 7 used a two-step process of inclusion of filamentary processes (Gu and Cheng 2015) to improve the results of several algorithms and were not included to provide a fair comparison among all base algorithms. Two variants were found too slow or did not generate results (PSF and LCMboost_2).We therefore considered only the remaining 35 base implementations (called “algorithms” hereafter) for bench testing on the *BigNeuron* image data (Supplementary Materials Table 2). The consensus tree algorithm introduced in this article was used to combine the results of all the algorithms.

### Algorithm bench-testing

To bench-test the algorithms, we ran all algorithms on the image stacks of gold standard reconstructions after pre-processing. Given that the testing of 35 algorithms in the Gold166 release image volumes (163 unique 3D stacks) implied running 5705 automatic reconstruction processes, those were parallelized in High-Performance Computing (HPC) facilities. The processing was distributed using the TITAN supercomputer at Oak Ridge National Laboratory (USA), as well as supercomputers at Lawrence Berkeley National Laboratory (USA) and Human Brain Project (Europe). Processes that took longer than 1 hour of computing time were terminated and did not provide results for the algorithm-dataset pair being tested. As a result, 5571 automatic reconstructions were generated (https://github.com/BigNeuron/Data/releases/tag/gold166_bt_v1.0). An example of a bench testing script for calling the tested algorithms can be found at: https://github.com/Vaa3D/vaa3d_tools/blob/d8e434c93708ab2a5bd349a79d9093d11aecf9d1/bigneuron_ported/benc h_testing/ornl/script_MPI/gen_bench_job_text_scripts_short.sh.

### Morphological analysis of neuronal reconstructions

To consistently compare morphological features that are dependent on the size of the trees, we first scaled both gold standard and automatic reconstructions using the pixel size information of each dataset (available at: https://github.com/lmanubens/BigNeuron/blob/main/scaling_gold.csv). Subsequently, to ensure consistent distance measurements in the 3D space and generation of consensus trees, we resampled the reconstruction nodes so that all the reconstruction segments had a length of 2 µm. To avoid loops, disconnected subtrees, and inconsistent hierarchies, we sorted the reconstructions using the “sort_neuron_swc” plugin (using the gold standard soma location as the root) in Vaa3D. Some of the tested algorithms did not perform radius estimation by default. In order to provide a comparable set of reconstructions, we estimated the radius of the trees using the “neuron_radius” plugin. Moreover, we analyzed the morphological features of post-processed trees with the “batch_compute” function of the “blast_neuron” plugin. Furthermore, we quantified the Sholl Intersection Profile scale, centripetal bias, and root angle distributions of the trees using the TREES toolbox functions “dissectSholl_tree” and “rootangle_tree” (Bird and Cuntz 2019).

### Interactive data analysis app

The datasets collected in this project are large, and the exploration is time-consuming. To allow fast interaction with the data, we developed an interactive analysis web app in Shiny (https://www.shinyapps.io/). Shiny allows the development of web applications that build on the R programming language utilities for statistical analysis and data plotting. Within the server, users can analyze dataset images, gold standard annotations, automatic reconstructions, and metadata associated with each dataset. The app allows users to interactively choose the image quality and tree morphology metrics used for dimensionality reduction and clustering analyses and perform reconstruction quality benchmarking. The Shiny app source code can be found at: https://github.com/lmanubens/BigNeuron/tree/main/shiny_app. A detailed description of the web-app organization and the analyses it performs can be found in the Supplementary Materials.

### Generation of consensus trees

Next, we developed an algorithm to iteratively merge the reconstructions obtained by different automatic algorithms. The aim of the algorithm is to conserve tree regions reliably retrieved by different algorithms and discard possible algorithm-dependent artifacts in order to obtain a consensus reconstruction that is potentially closer to the ground truth. A brief description of the steps of the algorithm can be found in the Supplementary Materials. An implementation of the algorithm can be found as a Vaa3D plugin called “consensus_skeleton_2” and the source code is publicly available at: https://github.com/Vaa3D/vaa3d_tools/tree/master/hackathon/xiaoxiaol/consensus_skeleton_2. We did statistical tests for errors in morphological metrics of automatic reconstructions using the “stat_compare_means” function of the ggpubr package (version 0.4.0).

### Prediction of the best automatic reconstruction algorithm

To predict the best algorithm within a set of automatic reconstruction methods, we used the neuromorphological features of the automatic reconstruction (Morphological analysis of neuronal reconstructions) and the image quality features of a dataset (Image quality measurement). The statistics were transformed using the Box-Cox method to ensure normality. We generated a support vector machine regression learner using the mlr3 (version 0.12.0) package in R (version 3.4.1). To generate learning curves, we used the “generateLearningCurveData” of the mlr (version 2.18.0). To generate the regression results of Fig. 6, the data was split by the ids of the datasets in 15% for testing and 85% for training sets. We predicted the value for the percentage of different structure between the automatic reconstruction and the gold standard with the regression model. We obtained the regression coefficient of determination with the “msr” function of the mlr3 package (version 0.12.0). We did Wilcoxon tests (two-sided) using the “stat_compare_means” function of the ggpubr package (version 0.4.0). The code for this analysis can be found at: https://github.com/lmanubens/BigNeuron/blob/main/mlr_regression/mlr3_regression_3DIQ_3.R.

### Showcase of best algorithm prediction on fMOST data

We predicted best-performing algorithms in fMOST datasets. 40 fMOST image volumes were kindly provided by Dr. Liya Ding (20 dendritic trees and 20 axonal trees). We processed the images with the same steps used in Gold166 dataset (see Section Image preprocessing), and obtained automatic and consensus reconstructions as described in Algorithm bench-testing and Generation of consensus trees. We correspondingly obtained image quality and neuromorphological features as described in Image quality measurement and Morphological analysis of neuronal reconstructions. After applying a Box-Cox transformation, all the features obtained were used as inputs to the SVM regression model obtained as described in the previous section.

## References

Alivisatos, A.P., Chun, M., Church, G.M., Greenspan, R.J., Roukes, M.L., and Yuste, R. (2012). The brain activity map project and the challenge of functional connectomics. Neuron 74, 970–974.

Bird, A.D., and Cuntz, H. (2019). Dissecting Sholl analysis into its functional components. Cell Rep. 27, 3081– 3096.e5.

Bray, M.-A., and Carpenter, A.E. (2018). Quality control for high-throughput imaging experiments using machine learning in Cellprofiler. Methods Mol. Biol. 1683, 89–112.

Cai, D., Cohen, K.B., Luo, T., Lichtman, J.W., and Sanes, J.R. (2013). Improved tools for the Brainbow toolbox. Nat. Methods 10, 540–547.

Capowski, J.J. (1983). An automatic neuron reconstruction system. J. Neurosci. Methods 8, 353–364.

Chiang, A.-S., Lin, C.-Y., Chuang, C.-C., Chang, H.-M., Hsieh, C.-H., Yeh, C.-W., Shih, C.-T., Wu, J.-J., Wang, G.-T., Chen, Y.-C., et al. (2011). Three-dimensional reconstruction of brain-wide wiring networks in Drosophila at single-cell resolution. Curr. Biol. 21, 1–11.

Chung, K., and Deisseroth, K. (2013). CLARITY for mapping the nervous system. Nat. Methods 10, 508–513.

Chung, K., Wallace, J., Kim, S.-Y., Kalyanasundaram, S., Andalman, A.S., Davidson, T.J., Mirzabekov, J.J., Zalocusky, K.A., Mattis, J., Denisin, A.K., et al. (2013). Structural and molecular interrogation of intact biological systems. Nature 497, 332–337.

Daigle, T.L., Madisen, L., Hage, T.A., Valley, M.T., Knoblich, U., Larsen, R.S., Takeno, M.M., Huang, L., Gu, H., Larsen, R., et al. (2018). A Suite of transgenic driver and reporter mouse lines with enhanced brain-cell-type targeting and tunctionality. Cell 174, 465–480.e22.

Ecker, J.R., Geschwind, D.H., Kriegstein, A.R., Ngai, J., Osten, P., Polioudakis, D., Regev, A., Sestan, N., Wickersham, I.R., and Zeng, H. (2017). The BRAIN Initiative Cell Census Consortium: lessons learned toward generating a comprehensive brain cell atlas. Neuron 96, 542–557.

Forbes, C., Evans, M., Hastings, N., and Peacock, B. (2011). Statistical distributions (Wiley Hoboken).

Gillette, T.A., Brown, K.M., and Ascoli, G.A. (2011). The DIADEM metric: comparing multiple reconstructions of the same neuron. Neuroinformatics 9, 233–245.

Gong, H., Xu, D., Yuan, J., Li, X., Guo, C., Peng, J., Li, Y., Schwarz, L.A., Li, A., Hu, B., et al. (2016). High-throughput dual-colour precision imaging for brain-wide connectome with cytoarchitectonic landmarks at the cellular level. Nat. Commun. 7, 12142.

Gu, L., and Cheng, L. (2015). Learning to boost filamentary structure segmentation. Proceedings of the IEEE International Conference on Computer Vision (ICCV) 639–647.

Gu, L., Zhang, X., You, S., Zhao, S., Liu, Z., and Harada, T. (2020). Semi-supervised learning in medical images through graph-embedded random forest. Front. Neuroinform. 14, 601829.

Hama, H., Kurokawa, H., Kawano, H., Ando, R., Shimogori, T., Noda, H., Fukami, K., Sakaue-Sawano, A., and Miyawaki, A. (2011). Scale: a chemical approach for fluorescence imaging and reconstruction of transparent mouse brain. Nat. Neurosci. 14, 1481–1488.

Huisken, J., Swoger, J., Del Bene, F., Wittbrodt, J., and Stelzer, E.H.K. (2004). Optical sectioning deep inside live embryos by selective plane illumination microscopy. Science 305, 1007–1009.

Jiang, S., Wang, Y., Liu, L., Ding, L., Ruan, Z., Dong, H.-W., Ascoli, G.A., Hawrylycz, M., Zeng, H., and Peng, H. (2022). Petabyte-scale multi-morphometry of single neurons for whole brains. Neuroinformatics 1–12.

Kanari, L., Dłotko, P., Scolamiero, M., Levi, R., Shillcock, J., Hess, K., and Markram, H. (2018). A topological representation of branching neuronal morphologies. Neuroinformatics 16, 3–13.

Kandel, E.R., Markram, H., Matthews, P.M., Yuste, R., and Koch, C. (2013). Neuroscience thinks big (and collaboratively). Nat. Rev. Neurosci. 14, 659–664.

Ke, M.-T., Nakai, Y., Fujimoto, S., Takayama, R., Yoshida, S., Kitajima, T.S., Sato, M., and Imai, T. (2016). Super-resolution mapping of neuronal circuitry with an index-optimized clearing agent. Cell Rep. 14, 2718–2732.

Li, A., Gong, H., Zhang, B., Wang, Q., Yan, C., Wu, J., Liu, Q., Zeng, S., and Luo, Q. (2010). Micro-optical sectioning tomography to obtain a high-resolution atlas of the mouse brain. Science 330, 1404–1408.

Li, R., Zeng, T., Peng, H., and Ji, S. (2017). Deep learning segmentation of optical microscopy images improves 3-D neuron reconstruction. IEEE Trans. Med. Imaging 36, 1533–1541.

Liu, S., Zhang, D., Song, Y., Peng, H., and Cai, W. (2018). Automated 3-D neuron tracing with precise branch erasing and confidence controlled back tracking. IEEE Trans. Med. Imaging 37, 2441–2452.

Markram, H. (2006). The blue brain project. Nat. Rev. Neurosci. 7, 153–160.

Meijering, E. (2010). Neuron tracing in perspective. Cytometry A 77, 693–704.

Meijering, E., Carpenter, A.E., Peng, H., Hamprecht, F.A., and Olivo-Marin, J.-C. (2016). Imagining the future of bioimage analysis. Nat. Biotechnol. 34, 1250–1255.

Meissner, G.W., Dorman, Z., Nern, A., Forster, K., Gibney, T., Jeter, J., Johnson, L., He, Y., Lee, K., Melton, B., et al. (2020). An image resource of subdivided Drosophila GAL4-driver expression patterns for neuron-level searches. bioRxiv 2020.05.29.080473

Nern, A., Pfeiffer, B.D., and Rubin, G.M. (2015). Optimized tools for multicolor stochastic labeling reveal diverse stereotyped cell arrangements in the fly visual system. Proc. Natl. Acad. Sci. U. S. A. 112, E2967–E2976.

Parekh, R., and Ascoli, G.A. (2013). Neuronal morphology goes digital: a research hub for cellular and system neuroscience. Neuron 77, 1017–1038.

Peng, H., Hawrylycz, M., Roskams, J., Hill, S., Spruston, N., Meijering, E., and Ascoli, G.A. (2015). BigNeuron: large-scale 3D neuron reconstruction from optical microscopy images. Neuron 87, 252–256.

Peng, H., Meijering, E., and Ascoli, G.A. (2015). From DIADEM to BigNeuron. Neuroinformatics 13, 259–260.

Peng, H., Zhou, Z., Meijering, E., Zhao, T., Ascoli, G.A., and Hawrylycz, M. (2017). Automatic tracing of ultra-volumes of neuronal images. Nat. Methods 14, 332–333.

Peng, H., Xie, P., Liu, L., Kuang, X., Wang, Y., Qu, L., Gong, H., Jiang, S., Li, A., Ruan, Z., et al. (2021). Morphological diversity of single neurons in molecularly defined cell types. Nature 598, 174–181.

Polavaram, S., Gillette, T.A., Parekh, R., and Ascoli, G.A. (2014). Statistical analysis and data mining of digital reconstructions of dendritic morphologies. Front. Neuroanat. 8, 138.

Radojević, M., and Meijering, E. (2019). Automated neuron reconstruction from 3D fluorescence microscopy images using sequential Monte Carlo estimation. Neuroinformatics 17, 423–442.

Santamaría-Pang, A., Hernandez-Herrera, P., Papadakis, M., Saggau, P., and Kakadiaris, I.A. (2015). Automatic morphological reconstruction of neurons from multiphoton and confocal microscopy images using 3D tubular models. Neuroinformatics 13, 297–320.

Senft, S.L. (2011). A brief history of neuronal reconstruction. Neuroinformatics 9, 119–128.

Shillcock, J.C., Hawrylycz, M., Hill, S., and Peng, H. (2016). Reconstructing the brain: from image stacks to neuron synthesis. Brain Informatics 3, 205–209.

Stockton, D.B., and Santamaria, F. (2017). Integrating the Allen Brain Institute cell types database into automated neuroscience workflow. Neuroinformatics 15, 333–342.

Treweek, J.B., Chan, K.Y., Flytzanis, N.C., Yang, B., Deverman, B.E., Greenbaum, A., Lignell, A., Xiao, C., Cai, L., Ladinsky, M.S., et al. (2015). Whole-body tissue stabilization and selective extractions via tissue-hydrogel hybrids for high-resolution intact circuit mapping and phenotyping. Nat. Protoc. 10, 1860–1896.

Verveer, P.J., Swoger, J., Pampaloni, F., Greger, K., Marcello, M., and Stelzer, E.H.K. (2007). High-resolution three-dimensional imaging of large specimens with light sheet–based microscopy. Nat. Methods 4, 311–313.

Wang, C.-W., Lee, Y.-C., Pradana, H., Zhou, Z., and Peng, H. (2017). Ensemble neuron tracer for 3D neuron reconstruction. Neuroinformatics 15, 185–198.

Wang, Y., Narayanaswamy, A., Tsai, C.-L., and Roysam, B. (2011). A broadly applicable 3-D neuron tracing method based on open-curve snake. Neuroinformatics 9, 193–217.

Winnubst, J., Bas, E., Ferreira, T.A., Wu, Z., Economo, M.N., Edson, P., Arthur, B.J., Bruns, C., Rokicki, K., Schauder, D., et al. (2019). Reconstruction of 1,000 projection neurons reveals new cell types and organization of long-range connectivity in the mouse brain. Cell 179, 268–281.e13.

Xiao, H., and Peng, H. (2013). APP2: automatic tracing of 3D neuron morphology based on hierarchical pruning of a gray-weighted image distance-tree. Bioinformatics 29, 1448–1454.

Zeng, H., and Sanes, J.R. (2017). Neuronal cell-type classification: challenges, opportunities and the path forward. Nat. Rev. Neurosci. 18, 530–546.

Zheng, T., Yang, Z., Li, A., Lv, X., Zhou, Z., Wang, X., Qi, X., Li, S., Luo, Q., Gong, H., et al. (2013). Visualization of brain circuits using two-photon fluorescence micro-optical sectioning tomography. Opt. Express 21, 9839–9850.

Zhong, Q., Li, A., Jin, R., Zhang, D., Li, X., Jia, X., Ding, Z., Luo, P., Zhou, C., Jiang, C., et al. (2021). High-definition imaging using line-illumination modulation microscopy. Nat. Methods 18, 309–315.

