## Supplementary Materials for "*BigNeuron*: A resource to benchmark and predict best-performing algorithms for automated reconstruction of neuronal morphology"

#### Image quality measurement

To quantify image quality, we implemented a plugin in Vaa3D (RRID:SCR_002609, version 3.497, http://vaa3d.org) that computes the features described in detail in (Bray and Carpenter, 2018). The following features are measured by the plugin.

##### *Blur*

- Focus score: a measure of the intensity variance across the image.

##### *Saturation*

- Percent maximal: percent of voxels at the maximum intensity value of the image.
- Percent minimal: percent of voxels at the minimum intensity value of the image.

##### *Intensity*

- Total intensity: sum of all voxel intensity values.
- Mean intensity, median intensity: mean and median of voxel intensity values.
- Std. intensity, MAD intensity: standard deviation and median absolute deviation (MAD) of voxel intensity values.
- Min. intensity, Max. intensity: minimum and maximum of voxel intensity values.

##### *Threshold*

- Otsu threshold: an automatically calculated threshold for each image that maximizes the inter-class variance between background and foreground (Otsu thresholding method; Otsu, 1979).

We also implemented Signal-to-Noise Ratio (SNR, Eq.1) and Contrast-to-Noise Ratio (CNR, Eq.2) Ratio quantifications, defining the boundary between foreground and background in the image volumes both as the average intensity and the Otsu threshold, respectively.

$$SNR = \frac{\overline{I_f}}{sd(I_b)} \quad \text{Eq. 1}$$

$$CNR = \frac{|max(I) - \bar{I}|}{sd(I_b)} \quad \text{Eq. 2}$$

Where $I_f$ and $I_b$ stand for the foreground and background intensity, respectively; $I$ stands for the image intensity; $max$ for the maximum value; and $sd$ for the standard deviation.

#### Image preprocessing

Due to varying properties of image acquisition pipelines among institutions, it is challenging to define a universal data preprocessing protocol for reconstruction. We performed the following preprocessing steps:

- MultiColor FlpOut (MCFO) color separation: For MCFO data, we applied the "neuron_color_separator" plugin of Vaa3D to extract single neurons and save each of them into a separate file for reconstruction. When more than one neuron had the same color, they we separated them with the connected component algorithm. Finally, we visually inspected all color separation results and filtered out cases in which more than one neuron were connected (i.e. had overlapping signals).
- Conversion to 8-bit single-channel images. We first simplified multiple color channel images to the main single channel. Subsequently, if the image had a 16-bit dynamic range, we converted it to 8-bit using linear rescaling.
- Color inversion: We inverted the intensity of bright-field images for reconstruction

For FlyCircuit and FlyLight datasets, bench testing automatic reconstructions were obtained both for the raw datasets, and after preprocessing with one of the two following possibilities:

- Gaussian smoothing: If the raw image had clear noise in the signal along the neurites, we used Gaussian smoothing with 7x7x5 Gaussian kernels. To do so, we ran the "image_filters/Gaussian_Filter" plugin in Vaa3D.
- Anisotropic filtering: It is able to enhance the signal along neurites while suppressing noise. This is useful for filling gaps when the intensity is uneven spatially. We did this using the "image_filter/anisotropic_filters" plugin in Vaa3D.

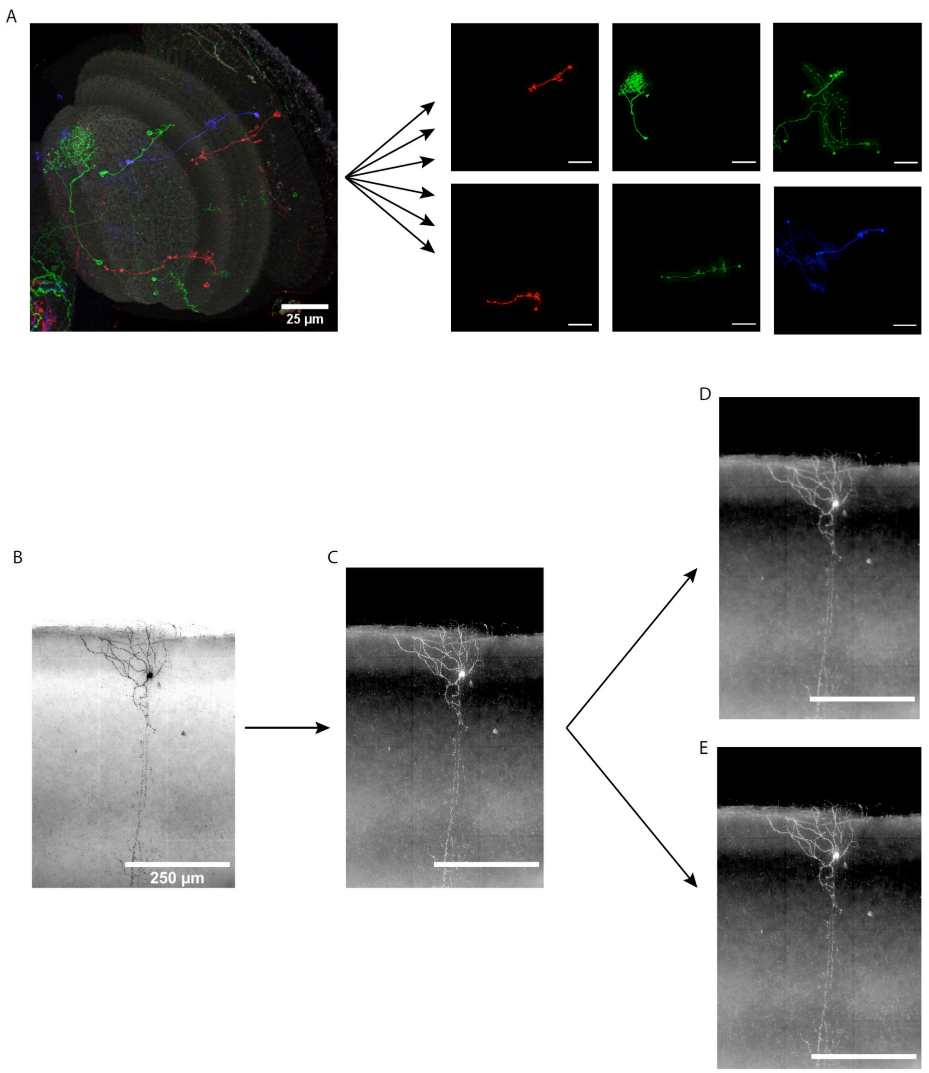


Supplementary Figure 1. Preprocessing steps used before automatic reconstruction algorithm bench testing.

**A** Schematic view of an example of color separation for a brainbow dataset. The image on the left shows a 2D maximum intensity projection of a brainbow RGB dataset. The six images on the right show separated individual neurons after using the “neuron_color_separator” plugin. The neurons labeled in the image are from the drosophila optic lobe from the Janelia Flylight dataset. **B** 2D maximum intensity projection of a brightfield microscopy dataset. Scale bar: 25 µm. **C** Shows the brightfield dataset after performing a color inversion. **D** Shows the inverted brightfield dataset after performing Gaussian smoothing. **E** Shows the inverted brightfield dataset after performing anisotropic filtering. The neuron in sections B-E is a pyramidal mouse neuron from barrel cortex provided by U.Goettingen. Scale bar: 250 µm.

#### Generation of gold standard reconstructions

We generated three files for each data entry: one 3D image stack file (in Vaa3D's .v3draw or .v3dpbd format), one neuron reconstruction file (in the .swc format), and one linker file (in Vaa3D's .ano format). Users can open any data entry conveniently by dragging and dropping the linker file (.ano) into the main window of Vaa3D. Since the original reconstructions validated by human annotators may not have consistent tags used to annotate dendritic, axonal, and somatic compartments, to avoid confusion and for simplicity, we uniformized all files so as include only two fixed node tags, 1 for the root and 3 otherwise.

#### Development of automatic reconstruction algorithms

All neuron tracing software is available from the latest binary release of Vaa3D. The reader is referred to the Vaa3D Github project page for more information (we documented the source code locations in https://github.com/BigNeuron/BigNeuron-Wiki/wiki/Neuron-Reconstruction-Algorithms). The tracing methods are available in the "main menu -> Plugins -> Neuron Tracing menu". Users can run these methods by using both GUI and the command line. Due to the limitation of dependency libraries for various tracing algorithms, the most complete set of tracing methods are available only for Linux.

#### Interactive data analysis app

##### *Organization of the web-app*

The web app has a set of panels that allow the users to select interactively data subsets based on their metadata, and metrics of interest. One left-sided panel allows users to choose the metrics of interest for dimensionality reduction and cluster analysis, and to choose which grouping variable is used for coloring in dimensionality reduction. A set of three bottom panels allows filtering the analyzed datasets by multiple selections of algorithms and datasets, and by filtering the range of values of a given metric. An additional bottom panel allows algorithm developers to upload a set of reconstructions obtained by self-developed algorithms to compare and benchmark new algorithms with the ones analyzed at the time of this publication.

##### *Dimensionality reduction*

To identify the metrics that better separate different groups of reconstructions, we performed a Scaled Principal Component Analysis (PCA, see Wold et al., 1987). We analyzed features associated with each reconstruction (morphological properties and quality metrics) and the datasets they belong to (image quality features). To obtain the results of Fig. 3, only gold standard reconstructions were analyzed, including the default set of morphology and image quality features defined in the Shiny app and summarized in Table 3. Before dimensionality reduction, the features with skewness greater than unity were log-transformed. We inverted negatively skewed distributions. This preprocessing ensures an approximately normal distribution of the input features (Polavaram et al., 2014). PCA was performed using "prcomp" in R (version 4.0.3). We generated biplots in R (version 3.4.1) using ggbiplot (version 0.55) to reduce the number of variables into a smaller number of principal components that account for most of the variance. Barplots of the contributions of each variable to the two first principal components were obtained using the factoextra package (version 1.0.7). To explore possible nonlinear relationships between datasets, we obtained t-distributed stochastic neighbor embedding (t-SNE, van der Maaten and Hinton, 2008) visualizations using the Rtsne package (version 0.15) with 500 maximum iterations and default parameters otherwise.

##### *Cluster analysis*

We obtained the pairwise Pearson correlation between every pair of features of all the neuron reconstructions and plotted them as a heatmap using heatmaply (version 1.1.0). We ordered the features in the heatmap based on hierarchical clustering and showed dendrograms on the top and right of the plot. We performed hierarchical clustering within the "heatmaply_cor" function, with the Ward.D2 method (Murtagh and Legendre, 2014). Similarly, we obtained dendrograms for hierarchical clustering of all the datasets using morphology metrics, image quality metrics, and the combination of both. To allow users to focus on specific groups of datasets, we defined clusters based on Gaussian Mixture model-based clustering results using the "Mclust" function of the mclust package (version 5.4.7). We obtained the dendrogram of figure 3I using the "hc" function of mclust. We calculated coefficients of determination and p values with the "stat_cor" function of the ggpubr package (version 0.4.0).

#### *Reconstruction quality benchmarking*

To measure differences between automatic and gold standard neuron reconstructions, we used the "neuron_distance" plugin in Vaa3D. This plugin quantifies the distance between neurons, defined as the average distance among all nearest point pairs. Given that the number of nodes can differ between pairs of reconstructions, distances are obtained twice using each reconstruction as a starting set for the search of nearest point pairs. Finally, the average bi-directional distance is calculated. Together with the average distance, the plugin also provides the percentage of nodes with pairwise distances higher than 2 voxels for each of the compared reconstructions (Peng et al., 2010). To assess reconstruction quality, we plotted the bi-directional average distance between pairs of neurons for each reconstruction method.

#### Generation of consensus trees

The consensus tree algorithm performs the following steps:

1.- K-Centroid clustering of all the nodes in input neurons. The number of clusters was defined as the average number of nodes of the input neurons.

2.- For each cluster resulting from the K-Centroid clustering, the center of the cluster was taken as a consensus node.

3.- Establish the weights among consensus nodes by collecting votes from the connections from individual input neuron trees. Every time a pair of nodes of two different clusters are connected in input trees, a vote is added to the connection between the consensus nodes of the respective clusters.

4.- Use a Minimum Spanning Tree algorithm (Prim, 1957) to connect consensus nodes to form the consensus tree.

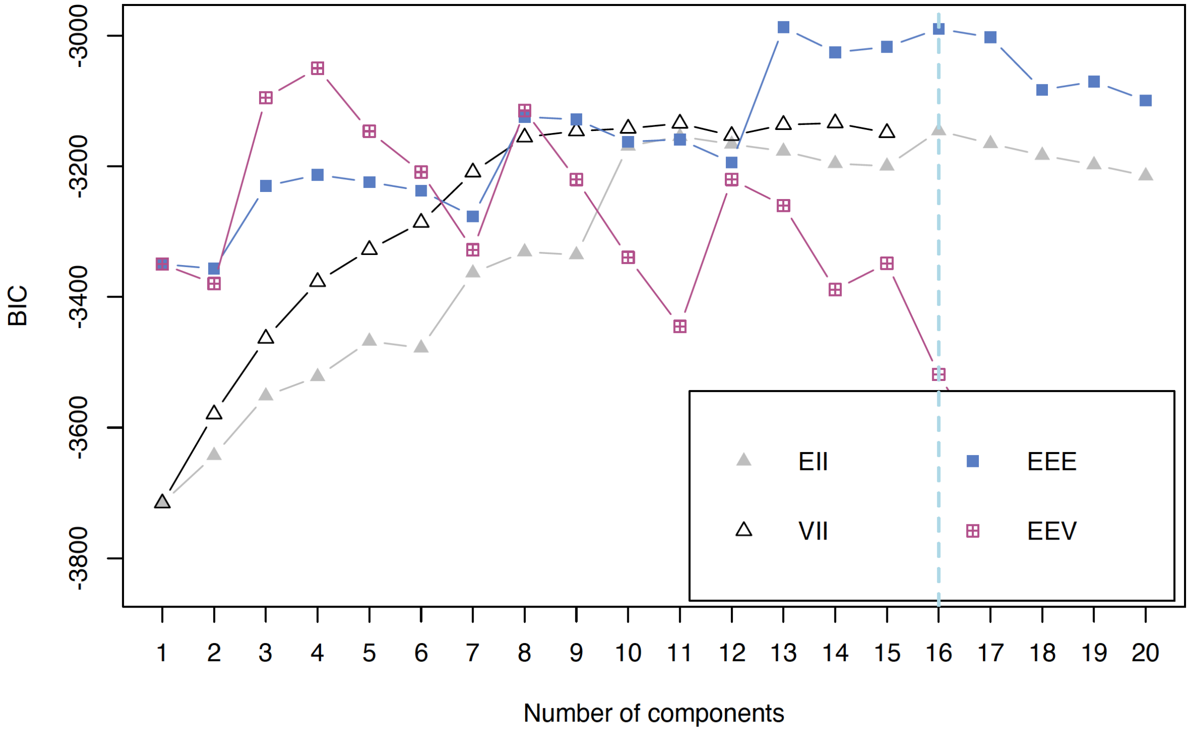


Supplementary Figure 2. Bayesian Information Criterion (BIC) for parametrized Gaussian Mixture models fitted by the expectation-maximization algorithm.

Each colored symbol indicates the BIC for a given mixture model with a number of components specified in the x axis. "EII": spherical, equal volume; "VII": spherical, unequal volume; "EEE": ellipsoidal, equal volume, shape, and orientation; "EEV": ellipsoidal, equal volume and equal shape. The dashed light blue line indicates the maximum BIC.

Table 1. Datasets included in the gold-standard Gold163.

| **Contact** | **Institution** | **Model organism** | **Age** | **Brain region** | **Cell type** | **Labeling method** | **Microscopy** | **Resolution** | **Reference?** | **DOI** | **In NeuroMorpho?** |
|---|---|---|---|---|---|---|---|---|---|---|---|
| G. Rubin | Janelia - Flylight | Fruit Fly | Adult | Many | Many | MultiColor FlpOut (MCFO) | Confocal | 0.38 isotropic | Yes | 10.1073/pnas.1506763112 and https://www.pnas.org/content/105/28/9715 | No |
| A. Chiang | Taiwan - Flycircuit | Fruit Fly | Adult | Many | Many | Gal4-GFP FLP-out labeling | Confocal | x-y:0.32 z:1 | Yes | 10.1016/j.cub.2010.11.056 | Yes |
| "Ryohei Kanzaki" and "Daisuke Miyamoto <>" | U Tokio | Fruit Fly | Adult | Many | Many | Immunostaining - anti-*Drosophila melanogaster* synaptotagmin | Confocal | Unknown | To database | https://invbrain.neuroinf.jp/modules/htmldocs/IVBPF/Top/index.html | No |
| Dan Cox & Giorgio Ascoli | GMU | Fruit Fly | Larvae | Cuticle, abdominal, segment 4-7 | Class I-IV | Gal4-UAS GFP transgenic labeling or F-actin GFP immunostaining | Confocal | Varies: x-y: 0.1483-0.2965-0.593 z: 0.5-1.5-2 | Unknown, has table with metadata: “Big Neuron-DA neuron dataset.xlsx” | 10.1007/s00429-017-1541-9 | Yes |
| "Ryohei Kanzaki" and "Daisuke Miyamoto <.i.u- | U Tokio | Silkmoth | 2–7 d after eclosion | Many | Many | Immunostaining - anti-*Drosophila melanogaster* synaptotagmin | Confocal | Varies x-y: 0.31-0.62-2 z: 0.70-1.44-2 | To database | https://www.ncbi.nlm.nih.gov/pmc/articles/PMC3431043/ | No |

| | | | | | | | | | | | |
|---|---|---|---|---|---|---|---|---|---|---|---|
| tokyo.ac.jp>" | | | | | | | | | | | |
| Rachel Wong | UW | Zebrafish | Larvae | Retina | RGC | *brn3c* MYFP transgenic expression | Confocal | Unknown | Yes | https://www.ncbi.nlm.nih.gov/pmc/articles/PMC1716713/ | No |
| Rachel Wong | UW | Zebrafish | Adult | Retina | RGC | *brn3c* MYFP transgenic expression | Confocal | x-y:0.21 z:25 | Yes | https://www.ncbi.nlm.nih.gov/pmc/articles/PMC1716713/ | No |
| Rachel Wong | UW | Zebrafish | Larvae | Retina | Horizontal | sws1:GFP transgenic expression | Confocal | x-y:0.07 z:0.25 | Yes | 10.1038/ncomms4699 | No |
| Rachel Wong | UW | Mouse | P24 | Retina | RGC | Anti-nonphosphone urofilament H , anti-ChAT , and anti-Lucifer Yellow | Confocal | Varies: x-y:0.1-0.33-0.35 z:0.3-1 | Yes | https://www.ncbi.nlm.nih.gov/pmc/articles/PMC3990865/ | |
| Jinhyun Kim | KIST | Mouse | Adult | Hippocampal | CA1 pyramidal | mGRASP | Confocal | x-y:0.104 z:0.5 | Yes | 10.1016/j.neuron.2013.11.026 | Yes |
| Jochen Staiger | UGoettingen | Mouse | P28 | barrel cortex, layer 2/3 | VIP | VIP-IRES-Cre::Ai9 transgenic | Confocal and Brightfield | x-y: 0.18310547-0.1606 z: 0.5522-0.5 | Yes | 10.1093/cercor/bhv202 | No |
| Manuel Peter | UCambridge | Mouse | 3 month | auditory cortex, layer 2/3 | excitatory neurons | Thy1.2-PA-GFP::NLS or R26 PA-GFP::NLS | 2-photon | x-y: 0.5377274 z:1 | Yes | 10.1371/journal.pone.0062132 | No |

| | | | | | | | | | | | |
|---|---|---|---|---|---|---|---|---|---|---|---|
| | | | | | | transgenic expression | | | | | |
| Yun Wang | Allen Institute | Mouse | P14–P21 | Primary visual cortex, layer II | Pyramidal | Single cell filling Biocytin | Confocal | x-y:0.3 z:0.5 | Yes | 10.1016/j.neulet.2015.02.043 | Yes |
| Hanchuan Peng / DeFelipe | Allen Institute | Human | 40yr | Cingulate cortex | Pyramidal | Single cell filling Biocytin | Confocal | x-y:0.24-0.31 z:0.42-1 | Yes | 10.1007/s40708-017-0063-9 10.3389/fnana.2013.00015 | Yes |
| Manuel Peter / R. Livesey | UCambridge | Human-cultured | 45 days after neuronal induction | NA | Excitatory neurons stem-cell derived | Synapsin GFP lentivirus and anti GFP staining | Confocal | Varies x-y: | Yes | 10.1038/nn.3041 | No |
| Ed Rubel and Yuan Wang | UW | Chicken | postnatal 4 to 10 days | nucleus laminaris in the auditory brainstem | Principal neurons | Single cell filling Alexa 488 | Confocal | Varies x-y:0.103-0.156 z:0.3- | Yes | 10.1002/cne.23520 | No |
| Hai-yan He Cline Lab | Scripps | Frog | st47-48 | tectum | Tectal neurons | Electroporation Gal4-eGFP | Confocal | x-y:0.01 z:1 | Yes | https://www.cell.com/neuron/pdfExtended/S0896-6273(16)30158-1 | |

Table 2. Reconstruction algorithms used in *BigNeuron*. The binary release v3.200 of Vaa3D contains functional binaries of the plugins. The source codes are available in the Vaa3D github repository. *BigNeuron* Table 2

| Name | Developed in E | Vaa3D plugin URL | Reference | Parameter sets | File name | Generated res |
|---|---|---|---|---|---|---|
| Advantra | Yes | https://github.com/Vaa3D/vaa3d_to | (Radojević and Meijering 2019) | Scale (scal)=10, Background ratio (bratio)=0.5, Correlation thresl | | Yes |
| Axis Analyzer | No | https://github.com/Vaa3D/vaa3d_to | (Arganda-Carreras et al. 2010 and Homan 2007) | Does not have parameters | | Yes |
| APP1 | No | https://github.com/Vaa3D/vaa3d_to | (Hanchuan Peng, Long, and Myers 2011) | inmarker_file=NULL, channel=0, bkg_thresh=40, If trace in | | Yes |
| | | | | inmarker_file=NULL, channel=0, bkg_thresh=10, If trace in | | Yes |
| | | | | inmarker_file=NULL, channel=0, bkg_thresh=AUTO, b_256 | | Yes |
| | | | | inmarker_file=NULL, channel=0, bkg_thresh=AUTO, b_256 | | Yes |
| APP2 | No | https://github.com/Vaa3D/vaa3d_to | (Xiao and Peng 2013) | inmarker_file=NULL, channel=0, bkg_thresh=10, b_256_cu | | Yes |
| Ensemble Neuron Tracer | Yes | https://github.com/Vaa3D/vaa3d_to | (C.-W. Wang et al. 2017) | Does not have parameters | | Yes |
| Ensemble Basic | Yes | https://github.com/Vaa3D/vaa3d_to | (C.-W. Wang et al. 2017) | Does not have parameters | | Yes |
| Ensemble V1 | Yes | https://github.com/Vaa3D/vaa3d_to | (C.-W. Wang et al. 2017) | Does not have parameters | | Yes |
| Ensemble V2N | Yes | https://github.com/Vaa3D/vaa3d_to | (C.-W. Wang et al. 2017) | Does not have parameters | | Yes |
| Ensemble V2S | Yes | https://github.com/Vaa3D/vaa3d_to | (C.-W. Wang et al. 2017) | Does not have parameters | | Yes |
| FarSight Snake | No | https://github.com/Vaa3D/vaa3d_to | (Narayanaswamy et al. 2011) | channel (starts from 1)=1 | | Yes |
| Fast Marching Spanning T | Yes | https://github.com/Vaa3D/vaa3d_to | (Wan et al. 2017) | Does not have parameters | | Yes |
| | | | | Multiscale Enhancement based (-f LCM_boost) | | Yes |
| | | | | Fast Marching based (-f LCM_boost_2) | | No |
| | | | | RegressionTubularityAC based (-f LCM_boost_3) | | Yes |
| | | | | mostVesselTracer based (-f LCM_boost_4) | | No |
| | | | | neuTube based (-f LCM_boost_5) | | No |
| | | | | SimpleTracing based (-f LCM_boost_6) | | No |
| | | | | APP2 based (-f LCM_boost_7) | | No |
| | | | | APP1 based (-f LCM_boost_8) | | No |
| | | | | FastMarching SpanningTree based (-f LCM_boost_9) | | No |
| LCMBoost | Yes | https://github.com/Vaa3D/vaa3d_to | (Gu and Cheng 2015) | NeuroGPS based (-f LCM_boost_10) | | No |
| Mean-shift | Yes | https://github.com/Vaa3D/vaa3d_to | (Wan et al. 2017) | channel=1, distance to delete covered nodes (prim_distan | | Yes |
| MOST ray tracer | No | https://github.com/Vaa3D/vaa3d_to | (Wu et al. 2014) | ch=1, threshold (th)=40, window size of the seed (seed)=2( | | Yes |
| MST: simple | Yes | https://github.com/Vaa3D/vaa3d_to | (Jian Yang et al. 2018) | channel=1, window size=5 | | Yes |
| nctuTW | No | https://github.com/Vaa3D/vaa3d_to | (Lee et al. 2012) | inmarker_file=NULL, threshold=0.9 | | Yes |
| nctuTW_GD | No | https://github.com/Vaa3D/vaa3d_to | (Lee et al. 2012) | Does not have parameters | | Yes |
| NeuronChaser (Miroslav) | Yes | https://github.com/Vaa3D/vaa3d_to | - | channel (starts from 1)=1, scale (scal)=10, Correlation thres | | Yes |
| NeuronGPS | Yes | https://github.com/Vaa3D/vaa3d_to | (Quan et al. 2016) | resolution=0.5 0.5 1, binarythreshold=15, tracevalue=10, e | | Yes |
| NeuroStalker | Yes | https://github.com/Vaa3D/vaa3d_to | - | channel=1, preprocessing=1, run unit-tests=1, step=5, step | | Yes |
| NeuTu autotrace | No | https://github.com/Vaa3D/vaa3d_to | (Jin et al. 2019) | Does not have parameters | | Yes |
| NeuTube | No | https://github.com/Vaa3D/vaa3d_to | (Zhao et al. 2011) | Does not have parameters | | Yes |
| PSF | No | https://github.com/Vaa3D/vaa3d_to | (Bas and Erdogmus 2011) | Currently unavailable for release due to extremely slow ru | | No |
| PYZH (Matlab) (Peng Yu | No | https://github.com/Vaa3D/vaa3d_to | - | min_soma_size=50, max_neurite_width=5, neurite_width | | Yes |
| RegMST (tubularity mode | Yes | https://github.com/Vaa3D/vaa3d_to | (Sironi et al. 2016; Hanchuan Peng, Ruan, Atasoy, et al. | $vaa3dv3.200ProgramPath/filter_banks/oof_fb_3d_scale_ | | Yes |
| Rivulet | Yes | https://github.com/Vaa3D/vaa3d_to | (Liu et al. 2016) | channel=1, threshold=2, quality=1, prune=10 | | Yes |
| SIGEN | No | https://github.com/Vaa3D/vaa3d_to | (Yamasaki et al. 2006; Minemoto et al. 2009) | Does not have parameters | | Yes |
| SimpleTracing: DFS | No | https://github.com/Vaa3D/vaa3d_to | (Jinzhu Yang, Gonzalez-Bellido, and Peng 2013) | Additional tracing method 2 (-f dfs) | | Yes |
| SimpleTracing: DT fields | No | https://github.com/Vaa3D/vaa3d_to | (Jinzhu Yang, Gonzalez-Bellido, and Peng 2013) | SimpleTracing (-f tracing) | | Yes |
| SimpleTracing: ray tracer | No | https://github.com/Vaa3D/vaa3d_to | (Jinzhu Yang, Gonzalez-Bellido, and Peng 2013) | Additional tracing method 1 (-f ray_shooting) | | Yes |
| SmartTracing | Yes | https://github.com/Vaa3D/vaa3d_to | (Chen et al. 2015) | Does not have parameters | | Yes |
| TReMAP | Yes | https://github.com/Vaa3D/vaa3d_to | (Zhou et al. 2016) | mip_plane=0, channel=1, bkg_thresh=10, If trace in a auto | | Yes |
| Consensus | Yes | https://github.com/Vaa3D/vaa3d_tools/tree/d71a3c4908871a1a63c69c60050b47207e1aeb( | | max_vote_threshold=3, clustering_distance_threshold=1( | | - |

Table 3. Summary of neuromorphologica, reconstruction quality and image quality features used, including a brief description and their units when applicable.

*BigNeuron* Table 3

| Feature set | Metric | Description | Unit |
|---|---|---|---|
| | Soma surface | Surface area of the soma, assuming it is spherical and using the rad | μm2 |
| | **Num. stems** | Number of branches stemming from the root | - |
| | Num. bifurcations | Number of bifurcations in the tree | - |
| | Num. branches | Number of segments in the tree between the root, branch points or | - |
| | **Num. tips** | Number of terminal points in the tree | - |
| | Overall x span | Unidimensional maximum distance between x coordinates of the rec | μm |
| | Overall y span | Unidimensional maximum distance between y coordinates of the rec | μm |
| | Overall z span | Unidimensional maximum distance between z coordinates of the rec | μm |
| | **Average diameter** | Average segment diameter | μm |
| | **Total length** | Sum of all the segment lengths | μm |
| Neuromorphological | Total surface | Sum of all the segment surfaces assuming each segment has a con | μm2 |
| | Total volume | Sum of all the segment volumes assuming each segment has a con | μm3 |
| | Max. euclidean distance | Maximum euclidean distance between any point of the reconstructio | μm |
| | **Max. path distance** | Maximum path distance along the tree nodes between any node of | μm |
| | **Max. branch order** | Maximum branch order, defined as 1 in the first branch stemming fro | - |
| | Average contraction | Contraction is defined as the ratio between euclidean distance and | - |
| | Average fragmentation | Average numer of segments in each branch | - |
| | Parent daughter ratio | Average ratio of the radius of parent and child segments in each bra | - |
| | Bifurcation angle local | Angle between downstream nodes closest to a bifurcation | degrees |
| | **Bifurcation angle remote** | Angle between downstream branch or termination points closest to a | degrees |
| | Average radius | Average segment radius | μm |
| | **Entire structure average from neuron 1 to 2** | Average distance among all nearest point pairs from bench-test reco | μm |
| | Entire structure average from neuron 2 to 1 | Average distance among all nearest point pairs from gold standard t | μm |
| | **Average of bidirectional entire structure average** | Average of entire strucuture average from neuron 1 to 2 and from ne | μm |
| Reconstruction quality | Different structure average | Average distane from neuron 1 to 2 and from neuron 2 to 1 for poin | μm |
| | Percent of different structure from neuron 1 to 2 | Percentage of nodes with pairwise distances higher than 2 micromet | - |
| | Percent of different structure from neuron 2 to 1 | Percentage of nodes with pairwise distances higher than 2 micromet | - |
| | **Percent of different structure** | Average of percent of different structure from neuron 1 to 2 and fror | - |
| | **Focus score** | A measure of the intensity variance across the image | - |
| | Percent maximal | Percent of voxels at the maximum intensity value of the image | - |
| | **Percent minimal** | Percent of voxels at the minimum intensity value of the image | - |
| | Total intensity | Sum of all voxel intensity values | - |
| | Mean intensity | Mean of voxel intensity values | - |
| | **Median intensity** | Median of voxel intensity values | - |
| | **Std intensity** | Standard deviation of voxel intensity values | - |
| | MAD intensity | Median absolute deviation (MAD) of voxel intensity values | - |
| Image quality | Min intensity | Minimum of voxel intensity values | - |
| | Max intensity | Maximum of voxel intensity values | - |
| | Otsu threshold | An automatically calculated threshold for each image that maximizes | - |
| | SNR mean | Signal-to-Noise Ratio (Eq. 1) defining the boundary between foregro | - |
| | CNR mean | Contrast-to-Noise Ratio (Eq. 2) defining the boundary between foreg | - |
| | SNR Otsu | Signal-to-Noise Ratio (Eq. 1) defining the boundary between foregro | - |
| | **CNR Otsu** | Contrast-to-Noise Ratio (Eq. 2) defining the boundary between foreg | - |
| | X-Y pixel size | Voxel size in the lateral dimension | μm |
| | Z pixel size | Voxel size in the axial dimension (along the optical axis) | μm |

### APPENDICES

#### Events (partial list)

1. [Oct 15, 2016] *BigNeuron* held a special session spanning a full day at the BIH'16, Omaha. In the morning 10 short talks were presented and in the afternoon 10 live demos were shown by the *BigNeuron* participants.

2. [May 28, 2016] *BigNeuron* held an exciting Big Neuroscience data workshop at Southeast University, China on May 27-28, 2016. 130+ people attended the meeting which featured a number of presentations from experts in China, Japan, Australia, Singapore, and USA.

3. [May 27, 2016] *BigNeuron* hackathon at University of Cambridge in 2015 has led to a newly accepted paper by Paulo Castro Aguiar group in the journal Neuroinformatics. This work has the title "N3DFix: An algorithm for automatic removal of swelling artifacts in neuronal reconstructions".

4. [Jan 2016] *BigNeuron* finished a successful visualization and analysis hackathon at Imperial College London's Data Science Institute (http://www3.imperial.ac.uk/newsandeventspggrp/imperialcollege/engineering/datascienceinstitute/eventssummary/event_4-12-2015-16-35-42). 20+ people from 7 countries attended this intensive event. Event was cosponsored by ICL, HBP, and Allen Institute.

5. [Nov 2015] *BigNeuron* held the first data visualization and analysis hackathon at Oak Ridge National Lab, utilizing the large display wall of supercomputing facility. (https://www.olcf.ornl.gov/2016/01/05/bigneuron-hackathon-branches-out-at-olcf/) . Event was cosponsored by INCF, ORNL, and Allen Institute.

6. [Oct 2015] A Neuron Tracing algorithm workshop devoted for *BigNeuron* was held together in the 2015 Bioimage Informatics conference at NIST (http://www.nist.gov/itl/ssd/is/bioimage-conference-2015.cfm). 5 teams from Janelia, HHMI (USA), Allen Inst. and Univ. of Georgia (USA), Taiwan, Singapore, and Illinois (USA), presented their featured work on various neruon tracing strategies.

7. [Aug 2015] The Big Machine for *BigNeuron* workshop, a supercomputing workshop featured *BigNeuron*, was held at London (http://braininformatics.london/)

8. [June 2015] *BigNeuron* data annotation workshop was held at Allen Institute. Experts of many different species gathered to reconstruct single neurons and generate gold and silver datasets for evaluating automated neuron tracing algorithms. (https://www.linkedin.com/pulse/bigneuron-seattle-neuron-annotation-workshop-june-15-17-hanchuan-peng)

9. [June 2015] *BigNeuron* held the third algorithm porting hackathon at Janelia Research Campus of HHMI, USA. 40 people attended this event! (First day: https://www.linkedin.com/pulse/bigneuron-janelia-hackathon-first-day-june-1-2015-hanchuan-peng and demo day: https://www.linkedin.com/pulse/bigneuron-janelia-hackathon-demo-day-june-5-2015-hanchuan-peng)

10. [May 2015] *BigNeuron* held the second algorithm porting hackathon at University of Cambridge, UK. Event was cosponsored by Wellcome Trust, University of Cambridge and Allen Institute for Brain Science. (https://www.linkedin.com/pulse/bigneuron-cambridge-hackathon-may-4-8-2015-hanchuan-peng)

11. [March 2015] The first *BigNeuron* algorithm porting hackathon kicked off at Beijing. NPR reported this event (http://www.npr.org/sections/health-shots/2015/03/31/396586398/hackers-needed-to-teach-computers-to-spot-sick-brain-cells). Event snapshots are here (https://www.linkedin.com/pulse/bigneuron-beijing-hackathon-hanchuan-peng)

#### Exemplar press releases

1. [Jan 2016] News about the *BigNeuron* Visualization and Data Analysis hackathon held at the Oak Ridge National Lab, Nov 2015. https://www.olcf.ornl.gov/2016/01/05/bigneuron-hackathon-branches-out-at-olcf/

2. [Sept 2015] The Brain Projects featured a *BigNeuron* page at http://brainprojects.onair.cc/category/data/bigneuron/ .

3. [May 2015] Oak Ridge National Lab issued a news piece about *BigNeuron* effort (https://www.ornl.gov/news/digitizing-neurons).

4. [May 2015] GEN published a news piece about *BigNeuron* (http://www.genengnews.com/gen-articles/brain-projects-get-researchers-thinking-big/5474/)

5. [April 2015] Lawrence Berkeley National Lab issued a news piece regarding its partnership in *BigNeuron* (http://cs.lbl.gov/news-media/news/2015/bigneuron-unlocking-the-secrets-of-the-human-brain/)

6. [April 2015] *BigNeuron* came to Cambridge! (http://www.neuroscience.cam.ac.uk/news/article.php?permalink=57dc178dfd)

7. [March 2015] GeekWire published its interview on *BigNeuron* (http://www.geekwire.com/2015/allen-institute-for-brain-science-leads-project-to-reconstruct-3d-neural-images-with-supercomputers/)

8. [March 2015] George Mason University issued a press release of the *BigNeuron* project (https://test.gmu.edu/news/1910)

9. [March 2015] The *BigNeuron* project goes public! News from Nature (http://www.nature.com/news/neuron-encyclopaedia-fires-up-to-reveal-brain-secrets-1.17232) , Science (http://www.sciencemag.org/news/2015/03/hacking-brain-one-cell-time), NBC (http://www.nbcnews.com/science/science-news/bigneuron-project-aims-untangle-brain-cell-structure-n334251), etc have nice coverage of this project.

10. [March 2015] Allen Institute issued the press release of the *BigNeuron* project (https://alleninstitute.org/press-release/international-initiative-launched-advance-state-art-digital-tracings-neurons)

#### Exemplar Publications (250+ related publications to date)

1. [March 31, 2017] The UltraTracer paper based on *BigNeuron* tools is published in Nature Methods (http://www.nature.com/nmeth/journal/v14/n4/full/nmeth.4233.html and http://biorxiv.org/content/early/2016/11/14/087726).

2. [March 30, 2017] The NeuronAssembler is published in Brain Informatics (https://link.springer.com/article/10.1007/s40708-017-0063-9).

3. [March 29, 2017] The M-MAST paper is published in BMC Bioinformatics by J. Wan and Ni. Zhong et al. (https://bmcbioinformatics.biomedcentral.com/articles/10.1186/s12859-017-1597-9)

4. [Feb 5, 2017] An ensemble neuron tracer by Wang et al, is published in Neuroinformatics (http://link.springer.com/article/10.1007/s12021-017-9325-1).

5. [Jan 30, 2017] The neuron tip detection paper by Liu et al was published in Pattern Recognition (http://www.sciencedirect.com/science/article/pii/S0031320317300511).

6. [Nov 14, 2016] The UltraTracer paper developed out of the *BigNeuron* project, was online at http://biorxiv.org/content/early/2016/11/14/087726 . This paper describes the first of its kind to trace/reconstruct arbitrarily large 3D image volume (e.g. with hundreds of billions of voxels) with any base tracer, effectively!

7. [April, 2016] *BigNeuron* project published a new paper on neuron reconstruction by Tom Cai's group from University of Sydney, the title of the work is "Rivulet: 3D neuron morphology tracing with iterative back-tracking" (http://link.springer.com/article/10.1007%2Fs12021-016-9302-0).

8. [Feb 24, 2016] The *BigNeuron* project published a new paper "Reconstructing the brain: from image stacks to neuron synthesis" (http://link.springer.com/article/10.1007%2Fs40708-016-0041-7), describing a very large scale generation and use of the *BigNeuron* data and tools.

9. [Aug 2015] The SmartTracing paper (also ported in *BigNeuron*) was published in Brain Informatics (http://link.springer.com/article/10.1007%2Fs40708-015-0018-y)

10. [July 2015] The Neuron journal published the position paper about *BigNeuron* project (http://www.cell.com/neuron/abstract/S0896-6273%2815%2900599-1)

11. [July 2015] The Neuroinformatics journal published an editorial on *BigNeuron* (http://link.springer.com/article/10.1007%2Fs12021-015-9270-9)

#### Data releases

1. [Nov, 2021] *BigNeuron* data are being hosted at Tencent AI – Southeast University collaboration website https://neuroxiv.net/bigneuron/ .

2. [Nov 2, 2016] The bench testing reconstructions of almost 8000 reconstructions for gold163 datasets have been released ( https://github.com/*BigNeuron*/Data/releases/tag/gold166_bt_v1.0 ).

3. [Feb 3, 2016] *BigNeuron* released the first set of 2000 single neuron image stacks for developing better tracing algorithms (https://github.com/BigNeuron/Data/releases/tag/data_v1.0_first2000). These datasets are fruitfly neurons and were originally from flycircuit database. The data have been standardized using the protocol (http://alleninstitute.org/bigneuron/data/) and shared with targeted developers in 2015. Now the datasets are completely open subject to the license terms at http://alleninstitute.org/bigneuron/participate/ .

4. [July 2015] First data release of *BigNeuron*: the training datasets of 79 neurons (https://github.com/BigNeuron/BigNeuron-Wiki/wiki/Bench-Testing-and-Training-Data#training-image-data) that come with gold-standard manual annotation have been distributed to algorithm developers to fine tune their methods.

#### Exemplar Presentations

1. [March 16, 2016] *BigNeuron* to be highlighted at APS'2016 Annual Meeting, Baltimore, USA in the session "Large Scale Neuroscience Projects". (http://meetings.aps.org/Meeting/MAR16/Session/B12)

2. [Dec 2015] First show of *BigNeuron* at Nanjing, China

(http://rcls.seu.edu.cn/newsread.aspx?newsid=446)

3. [Dec 2015] The *BigNeuron* project was introduced to Shanghai, China (http://www.shu.edu.cn/Default.aspx?tabid=10470&ctl=Detail&mid=19747&Id=89563&SkinSrc=[L]Skins/SDweb/index)

4. [Dec 2015] *BigNeuron* was highlighted in the South Korea KAIST neuroscience workshop (http://raphe.kaist.ac.kr/apctpkaist/schedule.htm)

5. [Nov 2015] AINI'2015 featured *BigNeuron* talk as a keynote (http://www.neuroinf.jp/aini2015).

6. [Sept 2015] The *BigNeuron* project was featured at Paris 2015 RDA Meeting on Infrastructure for Understanding the Human Brain (https://rd-alliance.org/plenary-meetings/sixth-plenary/programme/e-infrastructures-rda-data-intensive-science/infrastructure)

7. [Sept 2015] Hanchuan Peng highlighted the *BigNeuron* project in the CRCNS PI meeting at University of Washington. (https://crcns2015.wordpress.com/)

8. [Sept 2015] Giorgio Ascoli was featured for his talk at BIH'2015 and involvement of *BigNeuron* (http://braininformatics.london/tag/bigneuron/)

9. [Aug 2015] Erik Meijering gave a feature talk on *BigNeuron* at the 2015 EMBC conference at Milan, Italy (http://embc.embs.org/2015/w-fd-4/)

#### Open Data Agreement

The *BigNeuron* project imposes little restriction on the contribution, use, publication and distribution of the data, while providing as much flexibility as possible for contributors to continue their own work.

*BigNeuron* calls for the contribution of neuronal images for public domain bench testing. For any contributed neuron images, a set of reconstructions will be produced using the ported automated neuron tracing methods. The data contributors can use such reconstructions freely given the *BigNeuron* project is appropriately cited.

Contributors of such neuron images of *BigNeuron* project may use, copy, distribute, publicly perform, publicly display or create derivative works of the neuron reconstructions corresponding to respective images for research, noncommercial, and commercial purposes, given that *BigNeuron* project as well as the respective reconstructions and algorithms or implementations used in *BigNeuron* are appropriately referenced. In addition, where the reconstructions contain links to downloadable software applications, services or tools, image contributors may download and use such applications, services or tools as long as such contributors adhere to any license terms and conditions provided with those applications, services or tools.

Image contributors may not post reconstructions on social media or other third-party websites that require image contributors to acknowledge that they own the data they post (e.g., YouTube, Flickr and Twitter). Image contributors agree that they will not use the *BigNeuron* data in any manner that would violate anyone else's rights, such as copyright, trademark, patent, privacy or other rights. This includes removing any copyright, trademark or other proprietary notices from the reconstructions. Image contributors may not create hyperlinks to the *BigNeuron* resources (e.g. website, database, documentation site) that portray this project in a false or misleading light. Image contributors agree that they will only make lawful use of the *BigNeuron* Data and Resources in compliance with all federal, state, and local laws and regulations.

Image contributors may, and are encouraged to, develop new methods, applications, interfaces or other inventions or works that improve the use of, and build upon, the reconstructions. In order to make the reconstructions available to image contributors and others, however, the *BigNeuron* project must preserve its freedom to innovate. If image contributors develop improvements based on or utilizing the reconstructions, and such image contributors obtain any proprietary rights in or to that improvement, the image contributors and their successors or assigns agree that image contributors will not assert any claim for infringement against the *BigNeuron* project for the use of any improvement that was independently developed by or on behalf of the *BigNeuron* project. Additionally, the *BigNeuron* project retains its rights, title and interest in any reconstructions that are part of or are used by image contributors to create an Improvement.
